# A specialized mRNA translation circuit instated in pluripotency presets the competence for cardiogenesis in humans

**DOI:** 10.1101/2021.04.12.439420

**Authors:** Deniz Bartsch, Kaustubh Kalamkar, Gaurav Ahuja, Jan-Wilm Lackmann, Hisham Bazzi, Massimiliano Clamer, Sasha Mendjan, Argyris Papantonis, Leo Kurian

## Abstract

The blueprints for developing organs are preset at the early stages of embryogenesis. Transcriptional and epigenetic mechanisms are proposed to preset developmental trajectories. However, we reveal that the competence for future cardiac fate of human embryonic stem cells (hESCs) is preset in pluripotency by a specialized mRNA translation circuit controlled by RBPMS. RBPMS is recruited to active ribosomes in hESCs to control the translation of essential factors needed for cardiac commitment program, including WNT signaling. Consequently, RBPMS loss specifically and severely impedes cardiac mesoderm specification leading to patterning and morphogenesis defects in human cardiac organoids. Mechanistically, RBPMS specializes mRNA translation, selectively via 3’UTR binding and globally by promoting translation initiation. Accordingly, RBPMS loss causes translation initiation defects highlighted by aberrant retention of the EIF3 complex and depletion of EIF5A from mRNAs, thereby abrogating ribosome recruitment. We reveal how future fate trajectories are preprogrammed during embryogenesis by specialized mRNA translation.

**Teaser**: Cardiac fate competence is preprogrammed in pluripotency by specialized mRNA translation of factors initiating cardiogenesis

## Introduction

Embryonic development relies on precise and coordinated cell-fate decisions, a complex process that sculpts an entire organism from a single totipotent cell (Newport and Kirschner, 1982a, b; Spemann and Mangold, 2001; Xiang et al., 2020). The success of developmental cell-fate decisions requires timely, specific, accurate, and efficient rewiring of the regulatory proteome to support rapid cellular identity changes (Gao et al., 2017; Lu et al., 2009; van Hoof et al., 2012). In this regard, most research efforts over the last decades have concentrated on morphogen signaling, epigenetic, and transcriptional mechanisms (Gifford et al., 2013; Loh et al., 2016; Peng et al., 2019; Rogers and Schier, 2011; Stadhouders et al., 2019). However, selective translational control is arguably the primary determinant of regulatory protein abundance in mammals and thus proposed as a central regulator of embryonic cell fate decisions (Harnett et al., 2021; Kong and Lasko, 2012; Kristensen et al., 2013; Lu et al., 2009; Schwanhausser et al., 2011; Teixeira and Lehmann, 2019). Yet, how the developmental transcriptome is selectively translated to authorize cell fate decisions is a fundamental question that remains unclear.

The relevance of translational control in embryonic cell fate decisions is highlighted by the following evidence and concepts. While poorly understood, mRNA abundance does not reflect protein abundance at the systems level across evolution, especially during cell fate decisions (Buccitelli and Selbach, 2020; Lu et al., 2009; Vogel and Marcotte, 2012). Regulation at the level of translation would allow accurate and restricted subcellular abundance of fate regulatory proteins, thus enabling spatio-temporal precision in gene function without a need for *de novo* mRNA synthesis (Baser et al., 2019; Das et al., 2021). In addition, since translation is the most energy demanding cellular process, by decoupling mRNA abundance from protein abundance, selective translational control would ensure rapid and efficient responses during early embryogenesis, when energy availability is rate limiting (Hausser et al., 2019). In support, studies using mouse ES cells (mESCs) proposed an immediate and substantial increase in protein synthesis upon induction of differentiation, indicative of a systems-wide yet poorly understood reprogramming of the “translatome” (Ingolia et al., 2011; Sampath et al., 2008). *In vivo* studies in mice have suggested that, at the exit from pluripotency, the mesoderm lineage is particularly susceptible to translational control, while the regulatory mechanisms remain unclear (Fujii et al., 2017; Guzzi et al., 2018). Recently, an elegant proteomics-based study analyzing ribosome composition using tagged ribosomal subunits in mESCs suggested the role for proteins associated with ribosomal complexes (loosely termed “ribosome-associated proteins”, RAPs) in the selective control of translation while the role of RAPs in humans remains to be investigated (Simsek et al., 2017).

Taken together, despite the proposed prominent role of translational control in the developmental cell fate decisions, a systematic systems-wide understanding of the regulators, the molecular mechanism(s), and the principles by which the developmental transcriptome is differentially translated in time and space to allow cell-fate specification remains largely elusive, especially in humans (Shi and Barna, 2015). We address these outstanding questions using hESC-based cell fate decision models as a paradigm.

## Results

### ARC-MS identifies proteins recruited to translationally active ribosomes during mesoderm commitment

We hypothesized that the competence for embryonic cell fate decisions is translationally controlled by cell fate-specific RAPs, which control the selective and privileged translation of developmental regulators. However, the identification of RAPs recruited to ribosomes in a cell-fate specific manner is challenging due a number of reasons. Current methods used for identifying RAPs (i) either require the generation of engineered ribosomal proteins (Shi and Barna, 2015), (ii) suffer from contamination of unrelated protein complexes (Simsek et al., 2017) or (iii) cannot distinguish between translationally active and inert ribosomes (Ingolia et al., 2011; Sampath et al., 2008). During early embryogenesis, a significant fraction of ribosomes are inert and ribosomal proteins are generated in excess (Sampath et al., 2008). Therefore, prioritizing RAPs recruited to active ribosomes could be beneficial for identifying those that are functional in a cell fate-, context-, or stimuli-specific manner.

To cumulatively address these bottlenecks in the faithful identification of RAPs, we established Active Ribosome-Mass spectrometry (ARC-MS), a versatile and easy to implement method tailored for the identification of RAPs recruited to translationally active ribosomes (**Figure 1A**, detailed protocol in STAR **Methods**). Briefly, ARC-MS involves labeling of *de novo* synthesized proteins via a brief pulse (5 min) of cell-permeable, ‘clickable’ methionine analog (i.e., a derivative of non-canonical L-azidohomoalanine, AHA), followed by stably anchoring labeled nascent peptides to ribosomes using an anisomycin derivative (Minati et al., 2021). Active ribosomal complexes are isolated by ‘clicking’ nascent peptides harboring AHA directly to dibenzocyclooctyne beads. Next, RAPs along with ribosomal proteins and translation factors are biochemically separated from *de novo* synthesized nascent proteins (which remain covalently linked to the beads) and quantitatively detected independently via liquid chromatography coupled mass spectrometry (LC-MS/MS) (**Figure 1A**, *workflow*). Importantly, RAPs are identified by filtering out known ribosomal proteins, translation factors, and *de novo* synthesized proteins (detected independently after on bead digestion of peptides). Because of the short AHA pulse labeling, ARC-MS captures active ribosomal complexes at the early stages of translation, arguably the rate-limiting and most regulated stage of protein synthesis, further increasing the probability of identifying functional RAPs. The ability to experimentally separate *de novo* synthesized proteins from ribosomal complexes allows further stringency in the identification of RAPs.

**Fig. 1:**
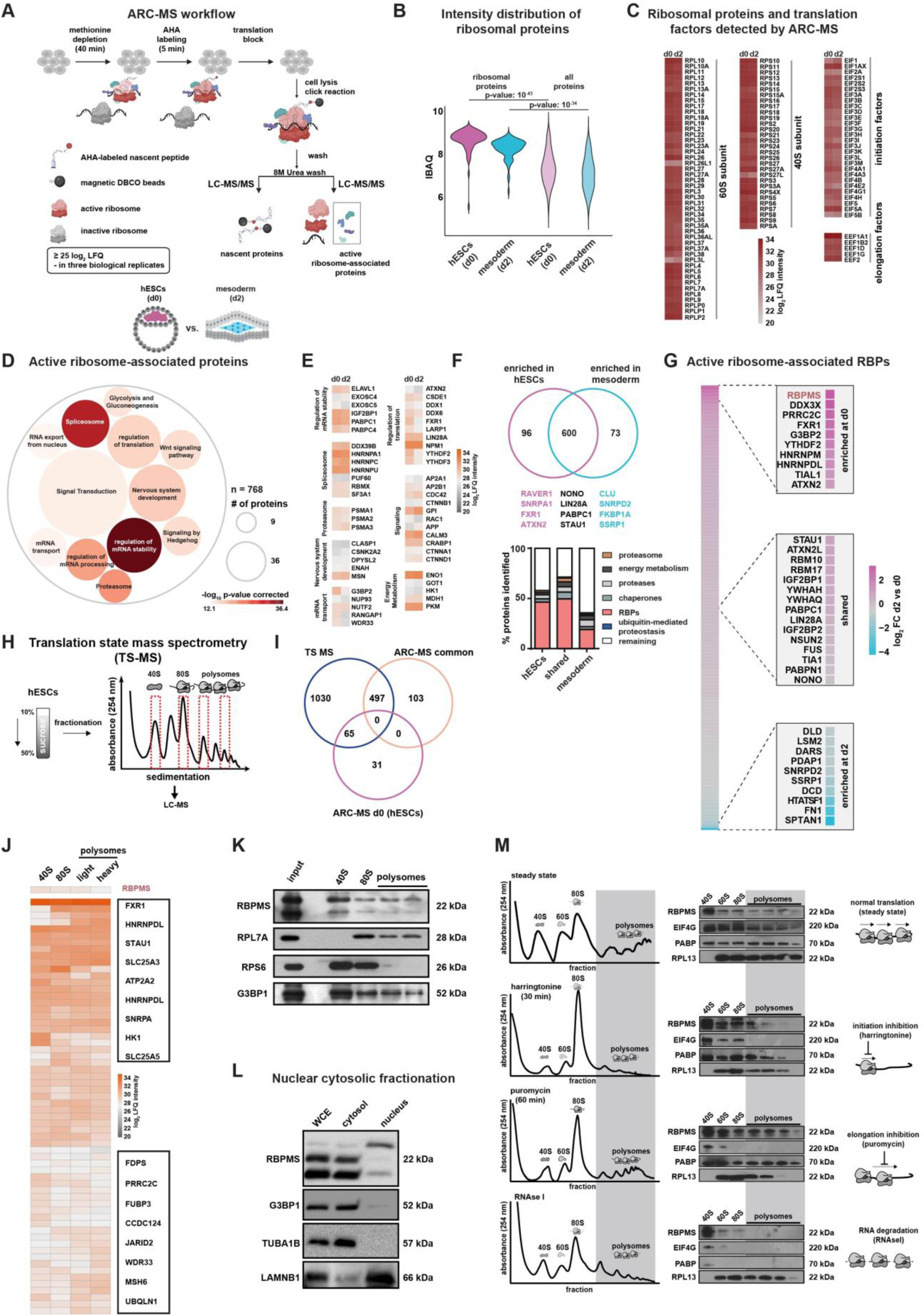
ARC-MS identifies proteins recruited to translationally active ribosomes during mesoderm commitment. **(A)** Schematic of ARC-MS workflow. ARC-MS was performed in hESCs (d0) and hESC-derived mesoderm progenitors (d2) (detailed protocol in **Methods**, biological replicates n=3). **(B)**, Violin plots depicting Intensity Based Absolute Quantification (IBAQ) values of ribosomal proteins and all identified proteins from ARC-MS data from hESCs Vs hESC derived mesoderm progenitors (only proteins detected in all three biological replicates above log_2_ LFQ ≥ 25, FDR ≤ 0.01 were considered). **(C)** Heatmap showing the enrichment of ribosomal proteins and translation factors (EIFs and EEFs) detected by ARC-MS in hESCs (d0) and mesoderm progenitors (d2). **(D)**, Gene Ontology-based functional enrichment analysis for proteins (excluding ribosomal proteins and translation factors) reliably identified by ARC-MS (shown are GO terms with p-value < 10^-12^). **(E)**, Heatmap depicting log_2_ label-free quantification (LFQ) enrichment of significantly enriched representative proteins recruited on active ribosomes from major GO term categories identified by ARC-MS. **(F)**, Venn diagram summarizing the distribution and enrichment (FC ±2, p-value ≤0.05) of proteins on active ribosomes in hESCs or mesoderm progenitors. Identified proteins categorized based on known molecular function depicted as percentage of total in bar graphs below in the indicated categories. **(G)** Heatmap displaying the enrichment of RBPs identified by ARC-MS between hESCs (d0) and mesoderm (d2). **(H)** Schematic outline of the underlying strategy employed for Translation state mass-spectrometry (TS-MS). (**I)** Overlap of proteins enriched at d0 and d2 ARC-MS with proteins detected via TS-MS in hESCs. **(J)** Distribution of proteins recruited on active ribosomes selectively in hESCs identified by ARC- MS) that are overlapping with TS-MS (n= 65) on indicated ribosomal fractions in hESCs. **(K)** Confirmation of RBPMS enrichment in ribosomal fractions by polysome profiling followed by immunoblotting. RPL7A, RPS6, and G3BP1 serve as controls. **(L)** RBPMS is predominantly a cytosolic protein, evaluated by western blot analysis upon nuclear/cytosolic fractionation, G3BP1, and TUBA cytosolic control, LAMINB1 nuclear control. **(M)** Residence of RBPMS on ribosomal complexes evaluated and confirmed by its characteristic association with the indicated complexes upon treatment with respective translation inhibitors, evaluated by polysome fractionation. Protein levels of RBPMS and the indicated controls on ribosomal fractions were detected by western blotting. Error bars represent ±SEM; p-values calculated using Student’s t-test, biological replicates n = 3.

As proof of principle for the systematic identification of RAPs regulating cell fate decisions, we selected the transition from pluripotency to mesoderm. We reasoned that the discord between active ribosomes and ribosomal abundance in pluripotency and the documented susceptibility of mesoderm lineage to translational control render this transition an ideal platform for identifying cell fate-regulating RAPs using ARC-MS. Therefore, we performed ARC-MS in hESCs (d0) and hESC-derived mesoderm (d2) (in 3 biological replicates/condition, **Figure S1A**) and only significantly detected proteins found in all three replicates (with log_2_ LFQ ≥ 25 in all replicates, FDR ≤ 0.01) were considered for further analysis (**Figure S1B** and **Table S1, Sheet 1** contains data for all proteins detected by ARC-MS in d0 and d2). To ensure successful and specific isolation of ribosomal complexes, we first calculated the intensity distribution of ribosomal proteins. They were the most abundant among the detected proteins (**Figure 1B**). Ribosomal proteins, translational initiation, and elongation factors constituted the majority of top enriched proteins, demonstrating successful isolation of active ribosomes (**Figure 1B, 1C**). The even distribution of ribosomal proteins on active ribosomes in both pluripotency and mesoderm indicated the faithful isolation of ribosomal complexes (**Figure 1C**). Notably, a few ribosomal protein isoforms showed differential enrichment, including RPL3L, -26L1 and -27L (Shi et al., 2017). A similar pattern was observed for translation factors also in support of recent studies illuminating their roles in selective translation.

Next, we filtered out ribosomal proteins and known translation factors in order to identify RAPs recruited on active ribosomes. In total, we identified 769 such RAPs in hESCs and hESC- derived mesoderm progenitors (**Table S1, Sheet 1**). Indicating direct crosstalk and synergy between the different stages of the mRNA life cycle and ribosomes, RAPs included proteins known to regulate pre-mRNA splicing, mRNA processing, stability, transport or export apart from known translation regulators (**Figure 1D** and **Table S1, Sheet 2** for details on all GO Terms). We identified proteins previously suggested to bind ribosomal complexes to regulate translation, such as FXR1, LIN28A, LARP1, ATXN1, DDX1, and PKM1 (Simsek et al., 2017) (a functionally characterized RAP in mESCs) (**Figure 1E**) (Shi and Barna, 2015). Notably, we identified various known regulators of embryonic development, energy metabolism, protein homeostasis, and components of morphogen signaling central to embryonic cell fate decisions on active ribosomes, including mediators of WNT signaling (**Figure 1E**). Previous reports using mESCs reported membrane proteins, centrosomes, clathrin complexes as well as the vault complex to be present as potential contaminants upon isolating ribosomal complexes to study associated proteins (Simsek et al., 2017). However, components of these complexes were scarcely present in ARC-MS data and were duly filtered out, although we cannot completely rule out transient interactions or shuttling of factors between complexes, which might be functionally relevant (**Figure S1G**). Thus, ARC-MS allows for the robust identification of RAPs recruited on actively translating ribosomes. Our hESC- to-mesoderm RAPs align with emerging hypotheses derived from bacteria, yeast, and mice that rather than constitutive protein synthesis factories, ribosomes can act as control hubs for cellular decision-making (Butland et al., 2005; Simsek et al., 2017; Vo et al., 2016).

### Cell fate-specific recruitment of RAPs on active ribosomal complexes revealed by ARC-MS

We identified 96 proteins to be preferentially recruited to active ribosomes in hESCs (fold change ≥2, p-value ≤0.05) as opposed to 73 in mesoderm and 600 were present in both states (**Figure 1F** and **Table S1, Sheet 1**). RNA binding proteins (RBPs) were the largest class of proteins (∼ 50%) among them. (**Figure 1F**, *bottom*). Considering the direct role of RBPs in the regulation of translation, we focused on RBPs recruited onto active ribosomes for further investigation (Shi and Barna, 2015). Among those shared between hESCs and mesoderm progenitors included known translational regulators, including LIN28A, IGF2BP1, STAU1, PABPC1, FUS, and TIA1 (Cho et al., 2012; Hentze et al., 2018; Tan et al., 2019). Mesoderm-specific ones were among the least known for their direct role in controlling selective translation, while it included RBPs such as LSM2, DARS and HTATSF1 (**Figure 1G**). Since we were interested in identifying regulators of selective translation controlling the transition from pluripotency to mesoderm, we focused on RAPs enriched on ribosomes in hESCs. The RNA binding protein RBPMS (<u>R</u>NA <u>B</u>inding <u>P</u>rotein with <u>M</u>ultiple <u>S</u>plicing) was among the top enriched RBPs recruited to active ribosomes in hESCs (d0) along with known regulators of selective translation, like FXR1, G3BP2, YTHDF2, TIAL1 and ATXN2. This consolidates that our identification and prioritization criteria yielded *bona fide* translational regulators (**Figure 1G** and **Table S1, Sheet 4**)(Hentze et al., 2018; Shi and Barna, 2015). RBPMS has never been reported as a regulator of translation, but has been proposed to be a potential regulator of embryogenesis, making it an ideal candidate for further investigation (Aguero et al., 2016; Farazi et al., 2014).

To independently verify the recruitment of the identified proteins to ribosomal complexes, we sought a global analysis of proteins that differentially associate with ribosomal complexes. To achieve this, we isolated ribosomal fractions from hESCs corresponding to 40S, 80S (monosome), and polysomes (light polysome fraction and heavy polysome fraction) by polysome profiling and subjected them to liquid chromatography coupled mass spectrometry (LC-MS) to identify their proteomic composition (translation state mass spectrometry, TS-MS) **(Figure 1H**, detailed protocol in Methods). The occurrence of the different ribosomal proteins in their expected fractions across all the samples indicated that the isolated fractions indeed represented the presumed ribosomal complexes (**Figure S1C**). After removal of ribosomal proteins and known translation factors, we identified 1408 proteins on ribosomal complexes, of which 600 were RBPs in line with our findings from ARC-MS (**Table S1, Sheet 6**). Similarly, proteins identified by TS-MS belonged to similar functional categories, e.g., RNA metabolic processes, mRNA processing, and translational control (**Figure S1D-S1F)**. Next, by intersecting the data from ARC-MS and TS-MS we could identify 66 proteins that are selectively recruited to ribosomal complexes in hESCs (**Figure 1I**). Importantly, RBPMS was also among the identified RBPs by TS-MS, thus confirming its recruitment on ribosomal complexes at the state of pluripotency (**Figure 1J**).

### RBPMS is preferentially recruited to active ribosomes in hESCs

Our candidate RAP in hESCs, RBPMS, is evolutionarily conserved across vertebrates and carries a single RNA recognition motif (Farazi et al., 2014; Teplova et al., 2016). While it has been suggested as a potential regulator of embryonic development based on studies in Xenopus and zebrafish, as well as binucleation of cardiomyocytes in mice; the function of RBPMS in human embryonic cell fate decisions, ribosome association, and translational control remain unknown (Aguero et al., 2016; Gan et al., 2022; Gerber et al., 2002). In line with a role in the regulation of translation, RBPMS sediments with ribosomal fractions and is enriched on the 40S ribosomal subunit (**Figure 1K)** similar to G3BP1, a regulator of selective translation known to associate with the 40S ribosomal subunit (Meyer et al., 2020). It is also predominantly cytosolic in hESCs (**Figure 1L**).

To validate the association of RBPMS with ribosomal complexes, we used orthogonal approaches. We reasoned that if RBPMS associates with translational machinery in hESCs, then upon treatment with specific translation inhibitors, it would show a characteristic shift in sedimentation commensurate to the inhibited step of translation. First, we used the specific translation initiation inhibitor harringtonine (2 µg/ml harringtonine for 30 min) and RBPMS was depleted from polysomes and concomitantly enriched in initiation fractions; changes in the enrichment of *bona fide* components of the translation machinery, EIF4G, PABP, and RPL13 serve as controls (**Figure 1M**, *top two panels*). We next treated hESCs with an inhibitor of elongation, puromycin (1 µg/ml for 1 h), to induce translational arrest. This led to the redistribution of RBPMS across fractions (**Figure 1M**, *third panel*). Then, we used RNAse I (5U, 30 min) treatment, which led to the disruption of ribosomal complexes and the accumulation of RBPMS in the 40S fraction (**Figure 1M**, *last panels*). This suggests a role of RBPMS in mediating translation initiation. To our knowledge, there are no other complexes that would show a similar sedimentation profile as ribosomal complexes and simultaneously show such characteristic change upon treatment with specific translation inhibitors. Considering its enrichment on the 40S complex in steady state and upon various treatments with translation inhibitors, notably upon RNAse I, we can rule out contamination by nascent RBPMS polypeptides. Thus, RBPMS is an active ribosome associated protein in hESCs.

### RBPMS loss causes global translation inhibition without affecting self-renewal in hESCs

To investigate the functional role of RBPMS, we generated a complete CRISPR/Cas9-mediated knockout in hESCs (hereafter RBPMS-KO) by targeting the exon-intron boundary of exon 1 with two gRNAs (**Figure 2A** and **Figure S2A**). Homozygous deletion of this exon-intron boundary disrupted the natural open reading frame of *RBPMS*, resulting in a complete loss of expression, which we confirmed both at the RNA and protein level (**Figure 2B** and **Figure S2B)**. Considering the recruitment of RBPMS on active ribosomes in hESCs, we next examined global translation in RBPMS-KO compared to WT using polysome profiling. We observed a severe reduction in the abundance and distribution of ribosomal complexes in RBPMS-KO hESCs where heavy polysomes (the most translationally-active fraction) were nearly absent (**Figure 2C**). This was followed by a 50% reduction in global *de novo* protein synthesis, reflected in the substantial reduction of newly synthesized proteins detected by short-term puromycin labeling evaluated either using an anti- puromycin antibody or fluorescent azide conjugated O-propargyl-puromycin (OPP) to avoid any detection biases (**Figure 2D and Figure S3A, 3B)**. Intriguingly, this severe global inhibition of translation upon loss of RBPMS did not alter the levels of pluripotency markers or self-renewal factors (**Figure S2C, 2D**) over a period of >20 passages. Critically, the global nascent transcriptional output of RBPMS-KO hESCs was only marginally affected (**Figure S3C)**. It also did not affect mitochondrial integrity based on MitoTracker^TM^ stained live-cell imaging (**Figure S3E**) or of co-staining for the mitochondrial marker MFN1 (**Figure S3D**), the cell cycle assessed by flow cytometry following PI staining (**Figure S3F**) and overall energy metabolism measured by Seahorse assay (**Figure S3G**). Together, these data reveal that loss of RBPMS inhibits mRNA translation without affecting the self-renewal of hESCs.

**Fig. 2:**
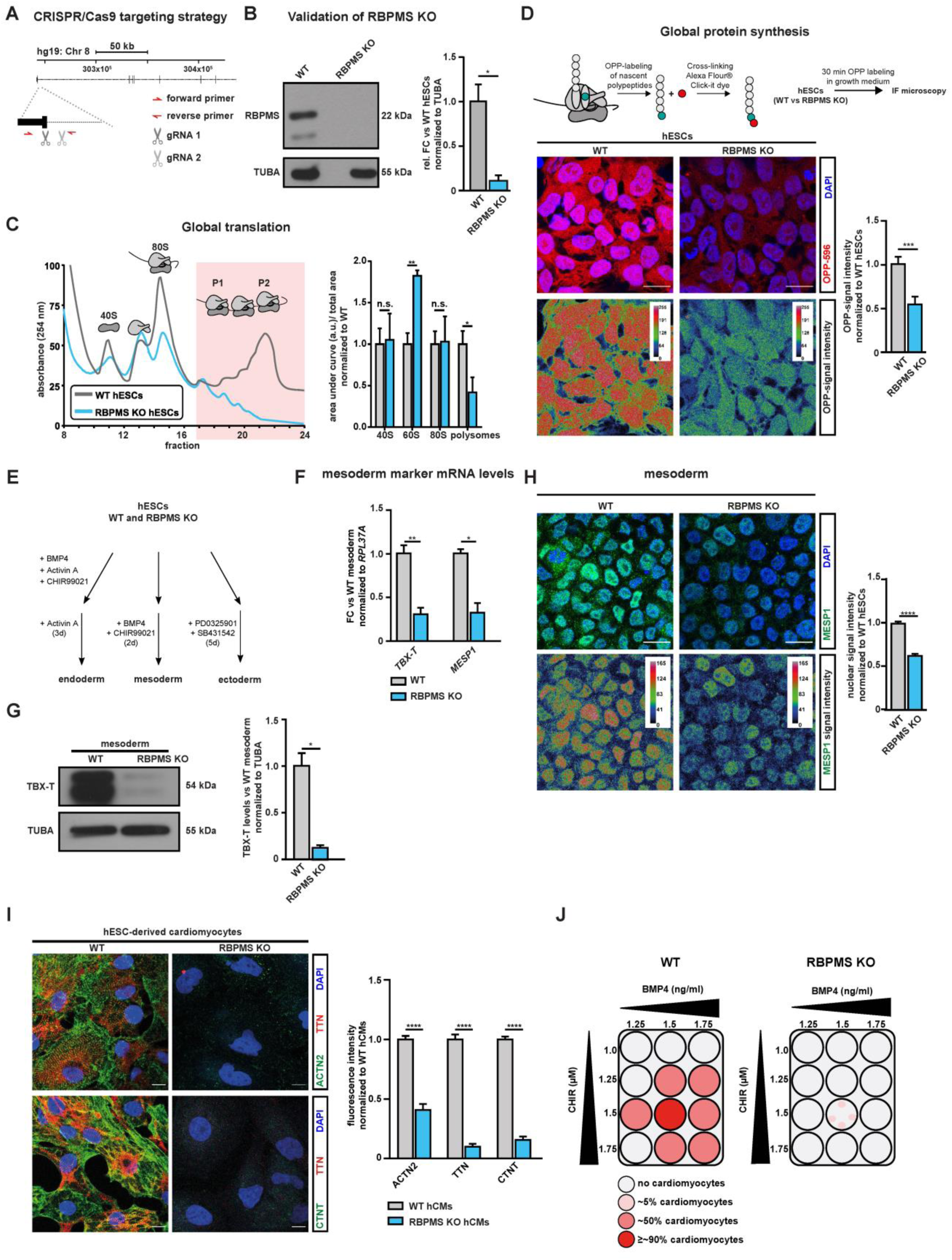
RBPMS loss causes global translation inhibition in hESCs without affecting self- renewal and specifically impedes cardiac mesoderm specification. **(A)** Schematic representation of RBPMS locus in humans and the CRISPR/Cas9-based targeting strategy used to generate homozygous RBPMS-KO **(B)** confirmed via western blot. **(C)** Loss of RBPMS impedes translation in hESCs, indicated by representative polysome profiles of RBPMS-KO hESCs w.r.t isogenic WT along with quantification of the area under the indicated ribosomal fractions on the right. **(D)** *De novo* protein synthesis is inhibited upon RBPMS loss, evaluated by measuring puromycin in cooperation on nascent proteins in RBPS-KO compared to WT by measuring uptake of OPP (quantifications on the right). **(E)** Schematic of lineage differentiation approaches used to determine the competence of RBPMS-KO hESCs to undergo germline commitment. **(F)** Mesoderm commitment is severely impaired upon loss of RBPMS as indicated by RT-qPCR for TBX-T (mesoderm) and MESP1 (cardiac mesoderm), as well as **(G)** western blot for TBX-T, quantification on the right. **(H)** Representative images of MESP1 staining upon mesoderm induction of RBPMS-KOs hESCs compared to WT, quantification on the right. **(I)** Immunofluorescence stainings for cardiac specific ACTN2 and TTN. Bar graph shows normalized expression levels of indicated cardiomyocyte markers. **(J)** Schematic summarizing the cardiac differentiation efficiency along the cardiac corridor for WT and RBPMS-KOs, indicating the inability of hESCs to terminally differentiate to cardiomnyocytes upon RBPMS loss. Error bars represent ±SEM; p-values calculated using Student’s t-test (*≤0.05, **≤0.01, ***≤0.001, ****≤0.0001, n = 3).

### RBPMS loss in pluripotency specifically impedes cardiac mesoderm specification

Because RBPMS loss abrogated translation homeostasis in hESCs, we reasoned that its loss would hamper cell-fate decisions enabling lineage commitment, a process heavily dependent on the *de novo* synthesis of fate commitment factors. To investigate the role of RBPMS in this process, we used a defined and directed differentiation method towards the three primary germ layers: ectoderm, mesoderm, and endoderm; recapitulating the early embryonic cell fate decisions (Frank et al., 2019) (**Figure 2E**).

Loss of RBPMS severely and specifically inhibited mesoderm commitment without affecting endoderm and ectoderm differentiation (**Figure 2F** and **Figure S4A-S4D)**. It abolished the ability of hESCs to effectively activate *TBX-T* (*BRACHYURY*), a master regulator of mesoderm commitment, and *MESP1,* a key early cardiac mesoderm marker (**Figure 2F-H**) (Bondue et al., 2008; Tosic et al., 2019). In addition, upon mesoderm induction, RBPMS-KO cells still expressed pluripotency factors aberrantly (**Figure S4F**) indicative of the inability of RBPMS-KO cells to efficiently exit pluripotency and undergo mesoderm lineage commitment upon instructive mesoderm morphogen signaling. To confirm that mesoderm commitment defects due to RBPMS loss were not a result of disrupted timing, we analyzed the expression dynamics of key mesoderm markers at close intervals. Markers such as *T* and *MIXL1* as well as WNT signaling mediators failed to activate in RBPMS-KO cells over the course of 24 h of mesoderm induction (**Figure S4E**)(Brade et al., 2013; Mazzotta et al., 2016; Rao et al., 2016).

Last, we tested whether the impaired differentiation of RBPMS-KO hESCs towards the mesodermal lineage detrimentally affects terminal fate choices. In this regard, we chose defined differentiation to cardiomyocytes since it is a robust, high-efficiency method allowing near- synchronous differentiation to a functional terminal fate (Frank et al., 2019; Rao et al., 2016). RBPMS-KO cells failed to properly activate key genes defining cardiac identity including aberrant expression of cardiac-specific transcription factors and sarcomeric genes and to produce cardiomyocytes in contrast to WT cells that constantly yielded homogenous populations of cardiomyocytes (**Figure 2I, Figure S4G**). Additionally, RBPMS-KO cells failed to produce cardiomyocytes across the “cardiac corridor” (**Figure 2J**), which is a BMP/WNT concentration grid for testing the ability of pluripotent stem cells to give rise to cardiomyocytes. Thus, we conclude that RBPMS is essential for accurate cell-fate decisions allowing mesoderm commitment and cardiac differentiation of hESCs.

### RBPMS is essential for cardiac mesoderm patterning and morphogenesis in human cardioids

Development of cardiomyocytes in the heart requires complex, rapid patterning and morphogenesis events in the cardiac mesoderm and the developing heart (events that cannot be recapitulated using directed 2D differentiation methods) (Brade et al., 2013). To understand whether loss of RBPMS in hESCs impedes cardiac mesoderm patterning, morphogenesis, and 3D organization (molecularly less understood processes in humans compared to cardiac cell fate decisions), we resorted to a recently introduced human cardiac organoid (cardioid) model (**Figure 3A** and **Figure S5A-S5C**) . Cardioids closely recapitulate otherwise hard to study cellular complexity, patterning and stratification of a developing human heart, including the molecular contributions from intercellular signaling and morphogenetic events, like chamber formation.

**Fig. 3:**
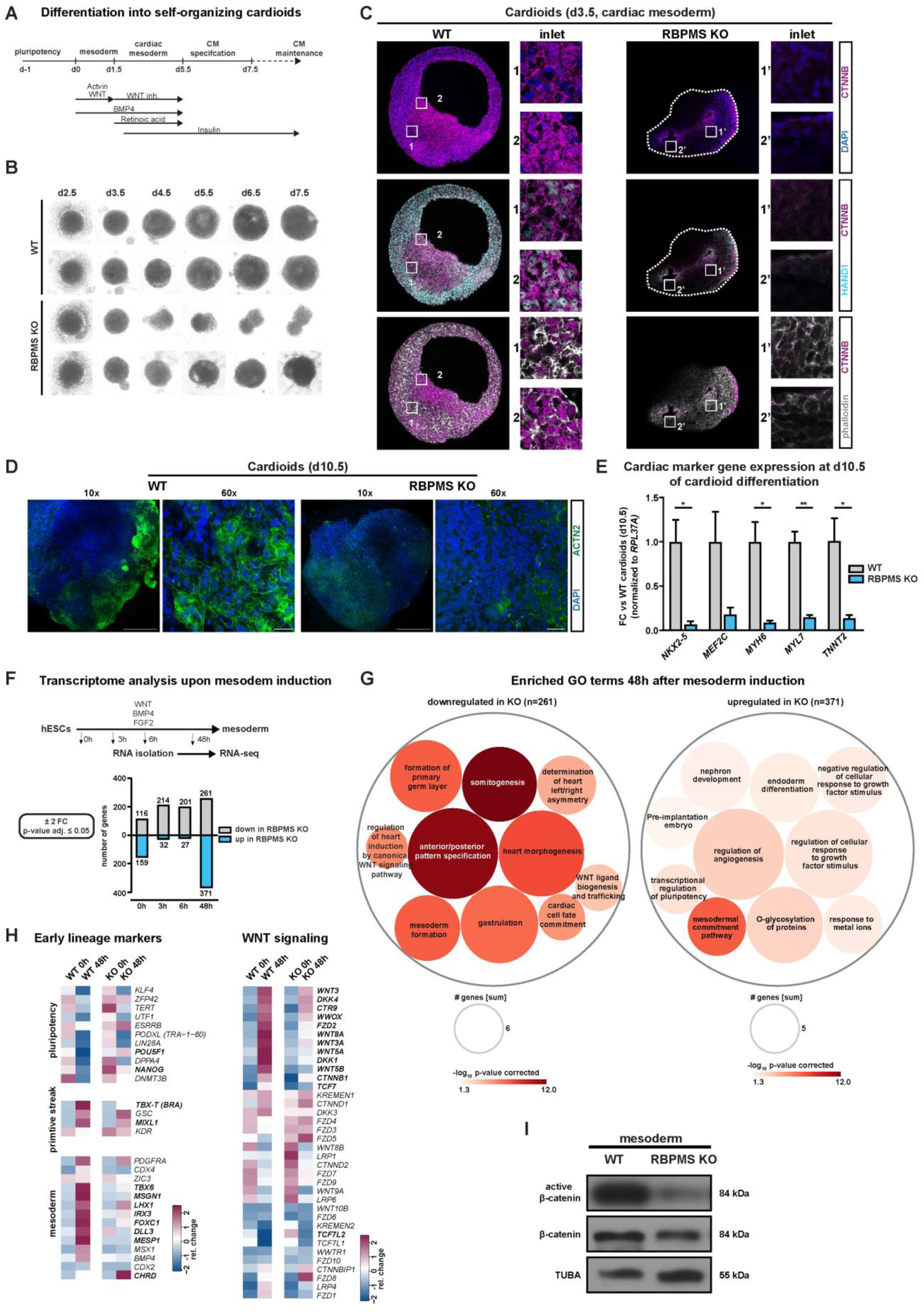
RBPMS is essential for cardiac mesoderm patterning and morphogenesis, revealed by human cardiac organoids (cardioids) **(A)** Schematic of cardioid generation. **(B)** Loss of RBPMS impairs cardioid formation at early stages of cardiogenesis as indicated by bright field images taken at indicated days during cardiac induction. **(C)** Whole organoid confocal imaging at cardiac mesoderm stage cardioids (d3.5) derived from RBPMS-KO and WT hESCs. Organoids were stained for cardiac mesoderm marker HAND1 and for evaluating the distribution of CTNNB and phalloidin to evaluate patterning and morphogenesis. Shown are stitched images from of whole organoid acquired by 63x objective. **(D)** Whole organoid imaging for cardiomyocytes specific ACTN2 in WT and RBPMS-KO Cardioids (d10.5). **(E)** RT-qPCR analysis of cardiac specific transcription factors and sarcomeric proteins performed on cardioids at d10.5. **(F)** Systematic identification of genes differentially expressed genes at indicated early and late time points during mesoderm induction upon RBPMS loss **(FC ±2, p-value ≤0.05, n=3)** evaluated by RNA-Seq, represented as bar graph. **(G)** GO-based analysis of genes downregulated and upregulated at 48h post mesoderm induction indicates defects in mesoderm and cardiac mesoderm cell fate specification and WNT signalling in RBPMS-KO cells w.r.t WT. **(H)** Heatmaps showing the expression of cell-fate markers representing the indicated stages of mesoderm induction between WT and RBPMS-KO. RBPMS-KO fail to silenence pluripotency markers, activate key primitive streak and mesoderm makers as well as WNT signaling components driving mesoderm induction. **(I)** Absence of RBPMS leads to reduced canonical WNT signaling activity as indicated by western blot for active beta catenin in mesoderm induced WT and RBPMS-KO hESCs. Error bars represent ±SEM; p-values calculated using Student’s t-test (*≤0.05, **≤0.01, ***≤0.001, ****≤0.0001, n = 3).

To investigate the effects on cardiac mesoderm patterning and cardiac morphogenesis by RBPMS loss in hESCs, we generated cardioids from RBPMS-KOs along a closely spaced time window between mesoderm commitment and cardiomyocyte specification (**Figure 3B**) and compared them to those derived from isogenic WT hESCs. RBPMS-KO cardioids were consistently smaller, lacking lumen and chambers, which often collapsed prematurely and disintegrated before reaching cardiac specification stages in line with their mesoderm commitment defects influencing patterning and morphogenesis (**Figure 3B, 3C**).

To further evaluate the role of RBPMS in cardiac mesoderm patterning, we analyzed the cardiac mesoderm-stage cardioids (d3.5). We focused on the distribution of cardiac mesodermal cells, lumen formation, and importantly the WNT-BMP signaling axis that is central to cardiac mesoderm patterning and morphogenesis (Brade et al., 2013; Hofbauer et al., 2021; Rao et al., 2016). First, WTs displayed a large lumen; RBPMS-KO often had multiple small lumens, which later did not fuse (**Figure 3C**). Second, indicating severe impairment in WNT signaling dynamics, the beta-catenin levels were low in RBPMS-KOs indicating severe impairment in WNT signaling dynamics (**Figure 3C**, *upper panel*). In WT d3.5 cardioids, the regions close to the periphery displayed mosaic distribution of cells in terms of nuclear/cell membrane localization of beta-catenin (**Figure 3C**, *upper panel inset 1*), while in the dense inner regions cells displayed uniform nuclear localization of beta-catenin (**Figure 3C**, *upper panel inset 2*). Beta-catenin was only present in small patches in RBPMS-KOs, and was near-exclusively localized to the cell membrane (**Figure 3C**, *upper panel insets 1’ and 2’*), indicating aberrant distribution and cardiac mesoderm patterning defects. In support of this, cells positive for HAND1 expression (a key WNT-BMP target marking cardiac mesoderm) were severely depleted in RBPMS-KO cardioids (**Figure 3C**, *middle panel insets 1, 2 and 1’, 2’*). The limited number of HAND1-positive cells in the RBPMS-KO scenario were distributed along the inner lumen compared to WTs. In WT cardioids, the inner part of the cardioids showed a mosaic distribution of HAND1-positive and -negative cells (**Figure 3C**, *middle panel inset 1*), while towards the periphery the cells were evenly HAND1-positive (**Figure 3C**, *middle panel inset 2*). However, such distinction was absent in RBPMS-KO with patches of HAND1-positive cells near the collapsed lumens (**Figure 3C**, *middle panel insets 1’ and 2’*), following the aberrant distribution and subcellular localization of beta-catenin. This points to severe cardiac mesoderm patterning defects in RBPMS-KOs. Importantly, in agreement with the inability of RBPMS-KOs to generate cardiomyocytes efficiently, the few RBPMS-KO cardioids that survived specification showed severe depletion of cardiomyocytes compared to corresponding WT (**Figure 3D**), which was accompanied by a near lack of expression of cardiac-specific transcription factors and sarcomere genes (**Figure 3E**). Together, our data reveal that RBPMS is central to cardiac mesoderm patterning and cardiomyocyte specification of hESCs.

### RBPMS is essential to activate the regulatory program instructing cardiac mesoderm

To carefully evaluate why RBPMS-KO fails to commit to cardiac mesoderm, we performed a whole-transcriptome analysis of RBPMS-KO compared to WT (n=3 biological replicates, poly(A)- selected mRNAs, paired-end 150-bp stranded libraries, ∼50 x 10^6^ clean reads/sample), in a closely- spaced window of mesoderm commitment differentiation (**Figure 3F** and **Table S2, Sheet 1**). The time points were chosen to evaluate the transcriptional differences at the stage of pluripotency (0h), upon receiving the mesoderm commitment cue (3h), early mesoderm (6h), and upon mesoderm commitment (48h).

First, at the transcriptome level, RBPMS-KOs and WTs were comparatively similar except for 275 differentially-expressed genes (log_2_FC ≥ (+/-) 1, p-value adjusted ≤ 0.05), arguing that the striking difference in the global translation upon RBPMS loss is not due to changes in mRNA levels. In addition, the genes differentially expressed did not have a direct connection to translational control or mesoderm/ cardiac mesoderm commitment (**Figure 3F**, *graph below* and **Table S2, Sheet 1**). We observed that most significant change in developmentally relevant gene expression signature at the 48h time point, aligning the cardiac mesoderm differentiation defect. Notably, Gene Ontology (GO) term analysis on differentially-expressed genes revealed that RBPMS-KO upon mesoderm induction were unable to activate the WNT signaling network, as well as those regulating gastrulation, mesoderm, cardiac cell fate commitment, and heart morphogenesis (**Figure 3G** and **Table S2, Sheet 2**). In contrast, RBPMS-KO cells retain aberrant pluripotency gene expression while also activating endodermal and ectodermal genes upon mesoderm induction (**Figure 3G**). Pluripotency factors, including *OCT4*, *NANOG*, *ESSRB*, and *ZFP42*, failed to be silenced in RBPMS-KOs even after 48 h, while primitive streak (e.g., *TBX-T, MIXL1*) and mesoderm markers (e.g., *MSGN1, LHX, MESP1, DLL3, TBX6*) were downregulated compared to WT hESCs (**Figure 3H**). Notably, RBPMS-KOs aberrantly activate early endodermal (e.g., *FOXA2, GDF3*) and ectodermal genes (e.g., *OTX2*) upon mesoderm induction suggesting that RBPMS loss disturbs germ layer decisions in hESCs (**Figure S5D**). Strikingly, a substantial number of genes involved in WNT signaling (e.g: *WNT3, WNT8A, WNT3A, WNT5B, CTNNB1, TCF7, DKK1, FZD2*) along with inhibitors of BMP signaling (e.g.: *CHRD*) were aberrantly expressed in RBPMS-KOs at the 48-h time-point (**Figure 3H** and **Figure S5D**). In RBPMS-KO hESCs, deregulation of WNT signaling components, which are key cardiac mesoderm specifiers, will lead to severe impairment of cardiac mesoderm specification, patterning, and subsequent cardiogenesis as evidenced in our 2D and cardioid models. This goes hand in hand with the reduced active beta- catenin levels in RBPMS-KO (**Figure 3I**). Thus, RBPMS is essential for hESCs to activate the gene regulatory network specifying cardiac mesoderm.

### mRNAs encoding cardiac mesoderm instructive gene regulatory network are targeted by RBPMS via 3’UTR binding

The repertoire of mRNAs targeted and regulated by RBPMS in hESCs is currently unknown. To comprehensively and stringently identify the network of mRNAs regulated by RBPMS in hESCs, we employed enhanced UV cross-linking and immunoprecipitation of ribonucleoprotein complex followed by massively parallel sequencing (eCLIP-seq) (**Figure 4A, Figure S6A and S6B**) (Van Nostrand et al., 2016). Following removal of PCR duplicates and normalization relative to size- matched input (SMI) controls from four independent replicates, we compiled transcriptome-wide, nucleotide-resolution, and high confidence binding maps displaying >80% overlap of target mRNAs between replicates (**Figure4B, Figure S6C-S6E** and **Table S3, Sheet 1**). Only statistically significantly enriched targets (fold enrichment over SMI ≥ 2, p-value ≤ 0.05) represented in all four replicates were considered for further analysis.

**Fig. 4:**
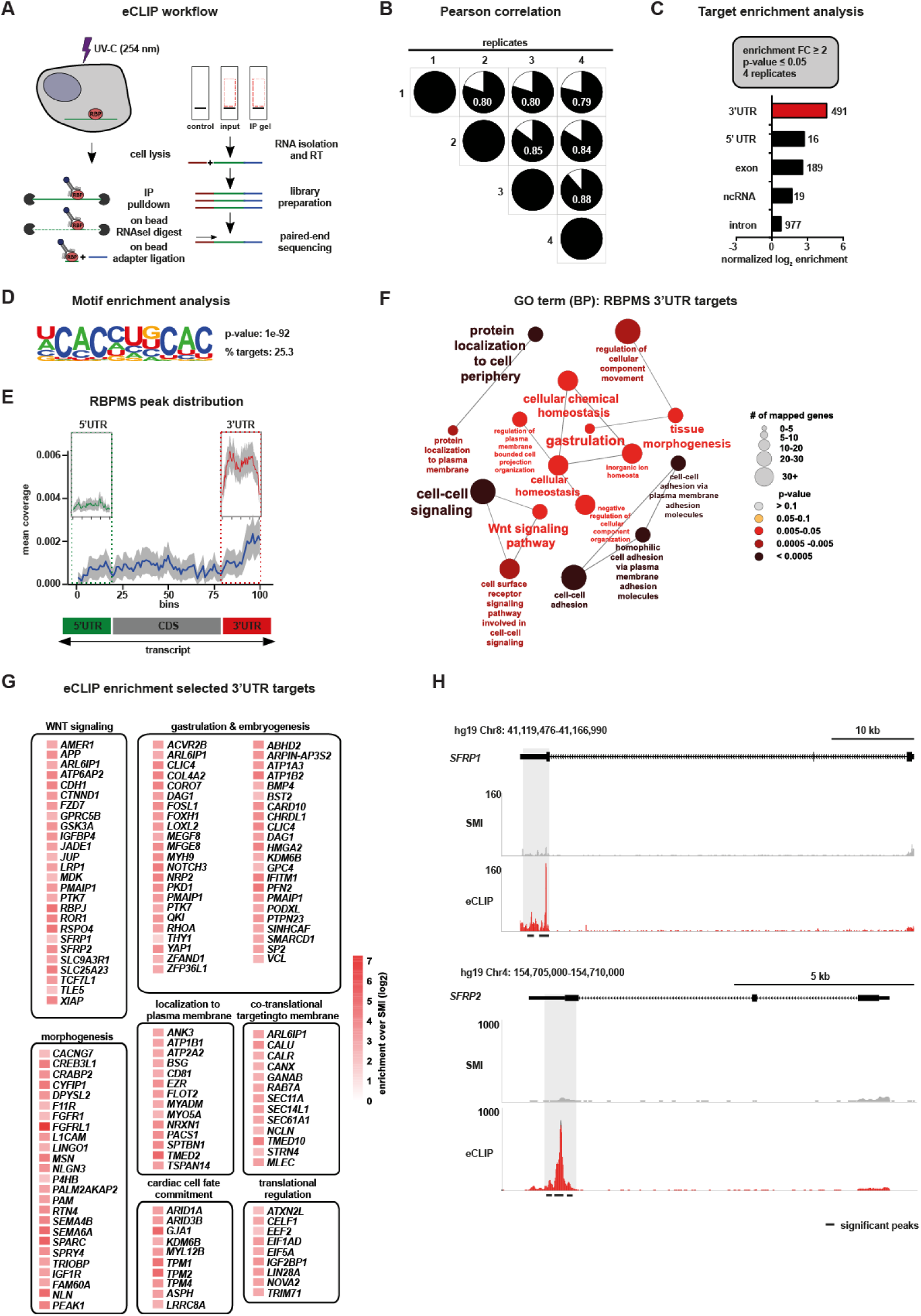
mRNAs encoding cardiac mesoderm instructive morphogen signaling are targeted by RBPMS via 3’UTR binding. **(A)** Schematic of the eCLIP-seq approach employed to faithfully generate a transcriptome-wide direct binding map for RBPMS at single-nucleotide resolution. **(B)** Biological quadruplicates of RBPMS eCLIP-seq show at least 80% overlap. Pie charts show the correlation of statistically significant uniquely mapped reads for each replicate over SMInput. **(C)** RBPMS reliably binds predominantly the 3’UTR of transcripts, demonstrated here by the distribution of the significantly enriched eCLIP peaks against the paired SMInput (fold change ≥ 2; p-value ≤ 0.05 in all 4 replicates). **(D)** Top sequence motif significantly bound by RBPMS. **(E)** Metagene plot visualizing the RBPMS peak distribution over SMInput illustrating prominent 3’UTR binding. **(F)** 3’UTR targets of RBPMS regulate molecular processes central to mesoderm/ cardiac commitment including WNT signal transduction, depicted by significantly enriched GO terms. **(G)** A curated set of RBPMS 3’UTR targets group based on their proven role in the indicated cellular, developmental and functional process, depicted as a heatmap of fold enrichment over SMInput. **(H)** Representative read density tracks show read density for RBPMS across the gene body of *SFRP1* and *SFRP2*, a representative target.

RBPMS was predominantly found to bind 3’UTRs of mRNAs (**Figure 4C** and **Table S3, Sheet 2**) on a bipartite CAC motif (**Figure 4D** and **Figure S6F**). RBPMS showed a higher degree of enrichment at the 3’UTR of target mRNAs compared to other regions, supported by normalized peak enrichment (**Figure 4C**). This was further evident when calculating RBPMS peak distribution in metagene plots, which showed substantial 3’UTR bias (**Figure 4E**), and in the representative loci of *SFRP1* and *-2* (**Figure 4H**).

Indicative of its direct role in regulating cardiac mesoderm fate of hESCs, the functional annotation of 3’UTR-bound mRNA targets revealed significant enrichment for key mediators of gastrulation, WNT signaling, as well as for cell surface and secreted proteins involved in developmental signaling (**Figure 4F** and **Table 3, Sheet 3**). A curated set of such 3’UTR targets, selected for high signal-over-input enrichment and grouped based on their molecular and developmental functions included genes from key categories involved in embryonic cell fate decisions. This included regulators of WNT signaling, e.g., *FZD7, GSK3A, SFRP1, SFRP2*; gastrulation and embryogenesis, e.g., *FOXH1, MYH9, YAP1, BMP4, SMARD1, SP2*; morphogenesis *FGFR1, MSN, IGF1R, NLN*; cardiac cell fate commitment/cardiac identity, e.g., *ARID1A, KDM6B, MYL12B, TPM1* (**Figure 4G**). In addition, mRNA encoding translational regulators were also bound by RBPMS at their 3’UTR, e.g., *CELF1, EEF2, EIF5A, IGFBP1 and LIN28A*, thereby offering an explanation of how its loss affects global translation in hESCs (**Figure 4G**). The 3’UTR targets of RBPMS encode proteins mostly localized to all subcellular locations, with bias for membrane and secreted proteins (**Figure S6G, Table S3, Sheet 4**). Together, we discovered that RBPMS directly targets a network of mRNAs encoding central regulators of early embryonic cell fate decisions, especially those critical for mesoderm instructive morphogen signaling, core components of the translation machinery, and regulators of mRNA translation.

### RBPMS controls the translation state of the regulatory components essential to initiate the cardiac mesoderm commitment in hESCs

To determine how the loss of RBPMS causes translation inhibition in hESCs and prevent cardiac mesoderm commitment, we first applied translation state RNA sequencing (TS-Seq) to investigate transcriptome-wide changes in the occupancy of ribosomal complexes upon RBPMS loss in hESCs. To this end, transcripts associated with ribosomal complexes (the 40S, 80S, light, and heavy polysomes) were isolated after ribosome fractionation, enriched for poly-adenylated transcripts, and subjected to transcriptome sequencing in parallel with total RNA from RBPMS-KO and isogenic WT hESCs (**Figure 5A**). To correct for technical variability and allow data normalization, two different sets of spike-ins were added to each fraction of our three biological replicates, after lysis and after polysome fractionation, respectively. We obtained 20-30 million clean reads per ribosomal fraction per replicate to ensure reliable quantification of ribosome occupancy differences in low to medium expressed transcripts.

**Fig. 5:**
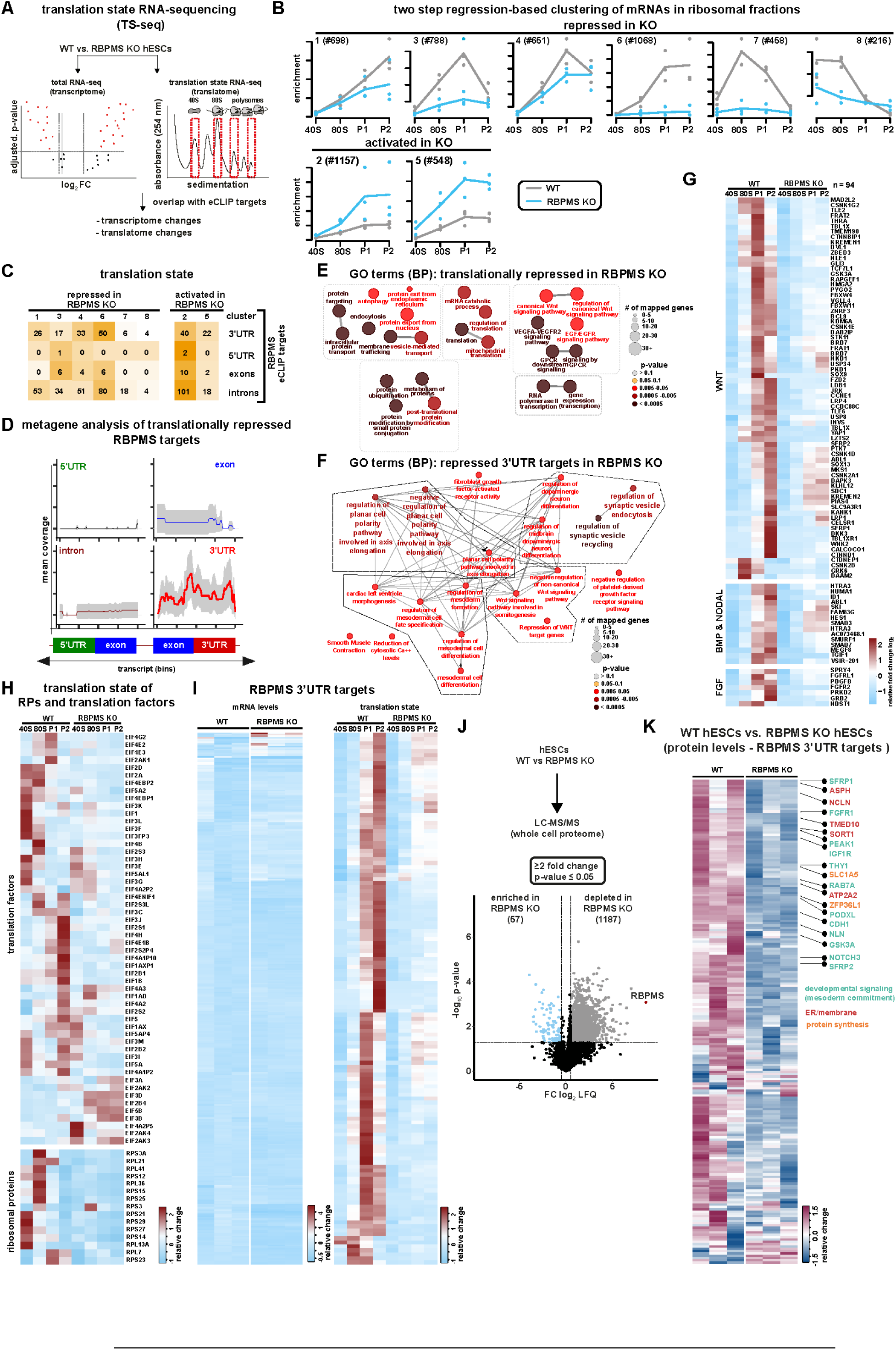
RBPMS controls the translation state of the regulatory components essential to initiate the cardiac mesoderm commitment in hESCs. **(A)** Schematic of the TS-seq strategy employed to evaluate the translational status of RBPMS-KO compared to WT (n=3). **(B)** Global impact of the loss of RBPMS on ribosome occupancy in hESCs, revealed by two-step regression analysis of the mRNAs enriching on indicated ribosomal fractions, derived from TS- Seq. **(C)** Translation state of mRNAs bound by RBPMS in RBPMS-KO compared to WT, grouped based on RBPMS binding coordinates, in the indicated translationally affected clusters identified by TS-Seq. **(D)** Metagene plot revealing RBPMS 3’UTR binding bias for translationally repressed RBPMS targets. **(E)** Functional analysis of all translationally repressed mRNAs and (**F)** translationally repressed 3’UTR targets in RBPMS-KOs vs WT hESCs illustrated as a significantly enriched curated list of GO terms. **(G)** Loss of RBPMS severely inhibits translation of the components vital mesoderm specifying signal transduction networks (WNT, BMP, NODAL, and FGF signaling), as well as **(H)** translation factors and ribosomal proteins. **(I)** mRNA bound by RBPMS at the 3’UTR are depleted from ribosomes in RBPMS-KOs without affecting the transcript levels. Heatmap on the left depicts mRNA levels of RBPMS 3’UTR targets while the heatmap on the right depicts their ribosome occupancy **(J)** Changes in total proteome between WT hESCs and RBPMS-KO hESCs depicted as volcano plot, derived from whole cell proteomics analysis. **(K)** Heatmap depicting protein abundance (as log2 LFQ) of RBPMS 3’UTR targets in WT and RBPMS-KO hESCs.

The loss of RBPMS resulted in severe translational inhibition (**Figure 2**), which makes the evaluation of changes in ribosome occupancy on specific mRNAs cumbersome. Therefore, a two- step regression-based clustering approach was used to identify meaningful differences in ribosome occupancy after normalization with dual spike-in controls. This approach allows for the identification of clusters of transcripts that were significantly different in their distribution of ribosomes, taking into account their occupancy across ribosomal complexes in the *RBPMS*-KO hESCs in comparison to isogenic WTs.

This way, we identified 8 mRNA clusters harboring >5500 mRNAs that exhibit a significant difference in their translation status following RBPMS-KO. Notably, upon RBPMS loss, ribosomal complexes were severely depleted in six clusters that harbored most of the translationally affected transcripts, while two clusters showed enrichment (**Figure 5B** and **Table S4**). Translationally repressed genes were crucial ones for cardiac cell-fate commitment, protein and mRNA metabolism (**Figure 5E)**, while translationally activated genes were involved in neurogenesis, endoderm and ectoderm development (**Figure S7B)** and **Table S4**). Integrative analysis of transcriptomics and TS-Seq data revealed that transcripts only showing transcriptional changes were not directly implicated in morphogen signaling or cardiac mesoderm development, the processes detrimentally affected by the loss of RBPMS (**Figure S7A** and **Table S4**), in agreement with our transcriptome analysis of RBPMS-KO versus WT hESCs (**Figure 3F** and **Table S2**).

Next, we investigated the translation status of RBPMS targets we identified in hESCs in relation to where it binds on the mRNAs (binding coordinates). In line with the prominent 3’UTR binding on developmentally relevant genes, substantial number of 3’UTR bound RBPMS targets were depleted from ribosomes in *RBPMS-KO* (**Figure 5C, 5I, Figure S7D Table S4**). Metagene analysis of translationally affected RBPMS targets revealed that the transcripts bound by RBPMS at the 3’UTR are depleted from ribosomes. In contrast, those bound at the 5’UTR, exons, and introns were not significantly affected. To further ensure that depletion of ribosomes from its 3’ UTR targets in the RBPMS-KO is not due to any indirect effects on mRNA levels, we systematically compared the transcript levels of 3’UTR targets and their ribosome occupancy upon loss of RBPMS. Substantial majority of the 3’UTR targets where translationally repressed in RBPMS-KOs (**Figure 5I, *heatmap on the right*)**, while the respective transcript levels of these targets remain largely unaffected (**Figure 5I, *heatmap on the left*)**. These data further reveal that RBPMS binding at the 3’UTR determines the translational status of its target mRNAs (**Figure 5D**).

Importantly, translationally inhibited 3’UTR targets of RBPMS encode regulators central to mesoderm specification, cell-fate commitment and, importantly, morphogen signaling, including WNT (**Figure 5F** and **Table S4**). Since morphogen signaling, particularly by WNT, BMP/NODAL, and FGF, defines mesoderm commitment from pluripotency, we then investigated the ribosome occupancy on mRNAs relevant to these processes (Brade et al., 2013; Hofbauer et al., 2020; Loh et al., 2016; Marvin et al., 2001; Steinhart and Angers, 2018). Strikingly, most WNT signal transduction components and those of BMP/NODAL and FGF signaling were severely depleted from active translational compartments upon RBPMS-KO (**Figure 5G** and **Table S4).** Off note, a subset of 3’UTR targets of RBPMS where translationally activated. They regulate processes not related to translation and cardiac mesoderm commitment (**Figure S7C**).

Surprisingly, we observed a depletion of mRNAs encoding core translation machinery, including translation initiation factors and ribosomal proteins in RBPMS-KO (**Figure 5H and Table S4)**. However, only a few of these were directly bound by RBPMS. This could further explain the global inhibition of translation upon loss of RBPMS (**Figure 2D, 2E, 5B**). This supports a model by which RBPMS selectively regulates the translation of its client mRNAs through 3’UTR binding and globally influences translation in an mRNA binding independent fashion. Its role in general translation could be through its direct interaction with translation machinery as indicated by ARC- MS and its association with 40S complex (**Figure 1G, 1M**).

RBPMS does not affect transcript stability (assessed for a selection of pluripotency factors and direct RBPMS 3’UTR targets involved in WNT signaling, following actinomycin-D treatment to inhibit transcription; **Figure S7E**).

RBPMS was suggested to regulate splicing in smooth muscle cells, extrapolated from targets identified by overexpression of RBPMS in HEK293T cells, which do not naturally express RBPMS (Nakagaki-Silva et al., 2019). A cursory analysis revealed minimal overlap between the targets reported in HEK293T cells and those we identified in hESCs, implying cell-type specificity. Nevertheless, despite the 3’UTR binding bias in RBPMS eCLIP data, we also detected low-affinity binding at introns (**Figure 4C, Table S3**)

To address whether RBPMS contributes to splicing in both hESCs (0h) and during mesoderm commitment (3h, 6h, and 48h post mesoderm induction), we computed splicing changes in RBPMS-KO. Our sequencing depth (∼50 Million clean reads/ sample/ replicate from stranded poly A selected library, n=3 biological replicates) allowed reliable investigation in changes in mRNA splicing(Van Nostrand et al., 2020; Yee et al., 2019). Briefly, we determined ψ scores for WT and RBPMS-KOs at the state of pluripotency and along mesoderm commitment time course, denoting significant changes with an FDR ≤ 0.01. A minimum of 10 junction reads were counted to compute inclusion/ exclusion events to robustly call splicing changes. Using a relaxed a cut-off of Δψ ≤ -0.5 (inclusion) or ≥ 0.5 (exclusion) to detect maximum number of splicing changes, we could only find minimal differences regardless of the time point investigated. Importantly, these few changes cannot explain the profound effects on mRNA translation or cardiac mesoderm commitment in RBPMS-KO cells (**Figure S8A, 8B** and **Table S5**). In fact, in d0 data, none of the 3’ and 5’UTR targets of RBPMS were affected at the level of splicing, while only 2 of its intronic and 1 of its exonic targets were among the genes whose splicing was affected. Similar trends were observed upon mesoderm induction time course, thus confirming our initial hypothesis that RBPMS controls mRNA translation without directly influencing mRNA splicing in hESCs and during mesoderm commitment (**Figure S8A, 8B** and **Table S5**).

Finally, we confirmed that the depletion of ribosomal complexes globally and from RBPMS 3’UTR targets results in a significant reduction in the abundance of corresponding proteins by performing an in-depth whole proteome analysis using LC-MS comparing WT and RBPMS-KO hESCs (**Figure 5J, 5K**). Notably, 1187 proteins where depleted in RBPMS-KO while on 57 where enriched confirming that RBPMS is essential for protein homeostasis in hESCs. Specifically, the protein levels of the 3’UTR targets of RBPMS where significantly depleted in RBPMS-KOs including WNT signal transduction components and mesoderm regulators (**Figure 5K, Figure S7F**), thus confirming the pivotal role of RBPMS in controlling their protein abundance in hESCs. Together, our data reveal that RBPMS is key regulator of mRNA translation in hESCs, primarily controlling the components of mesoderm instructive morphogen signaling, cell fate decisions and mRNA translation (summarized by the translational status of the curated list of key genes affected by its loss, including key 3’UTR targets, **Figure 5J**). Thus, RBPMS primes the selective translation of factors essential to initiate cardiac fate programming in hESCs, revealing that the competence for committing to the cardiac fate is predetermined by RBPMS-mediated translation circuit already instated in pluripotency.

### RBPMS specializes mRNA translation in pluripotency, selectively via 3’UTR binding and globally by controlling translation initiation and ribosome recruitment

Our data so far suggest that the role of RBPMS in translation could be two-pronged (i) as an activator of selective translation of mesoderm instructive cell-fate regulators via 3’UTR binding, and (ii) as a general regulator of translation in hESCs via recruitment to ribosomal complexes.

Therefore, we asked whether RBPMS binding at the 3’UTR can selectively control translation in hESCs. To this end, we first generated a destabilized dsRED based-reporter system carrying the 3’UTRs of two RBPMS targets *SFRP1* and *SFRP2* (inferred by RBPMS eCLIP-Seq in hESCs) (**Figure 6A**). The translation state and protein levels of both *SFRP1* and *SFRP2* are RBPMS-dependent in hESCs (**Figure S7D, S7G**). As evidenced by time-lapse microscopy, dsRED signal intensity was severely reduced in RBPMS-KO compared to WT hESCs (**Figure 6B**). This reduction could be rescued by ectopically expressing wild-type RBPMS, but not by RBPMS mutant carrying a point mutation (K100E) abolishing the RNA binding ability (**Figure 6B**). To confirm the ability of RBPMS to selectively control translation in a 3’UTR-dependent manner, we generated a set of dual luciferase-based bicistronic reporter constructs carrying with or without RBPMS binding sites (with *SFRP1* or *ACTB* 3’UTR, respectively) (**Figure 6C**, *illustration*). RBPMS loss led to significant reduction in luciferase activity for *SFRP1*-3’UTR fusions while *ACTB-3’UTR* and luciferase-only controls remained unaffected; **Figure 6C**). Collectively, these reporter based assays reveal that, RBPMS selectively activates translation of mRNAs carrying its binding sites at the 3’UTR.

**Fig. 6:**
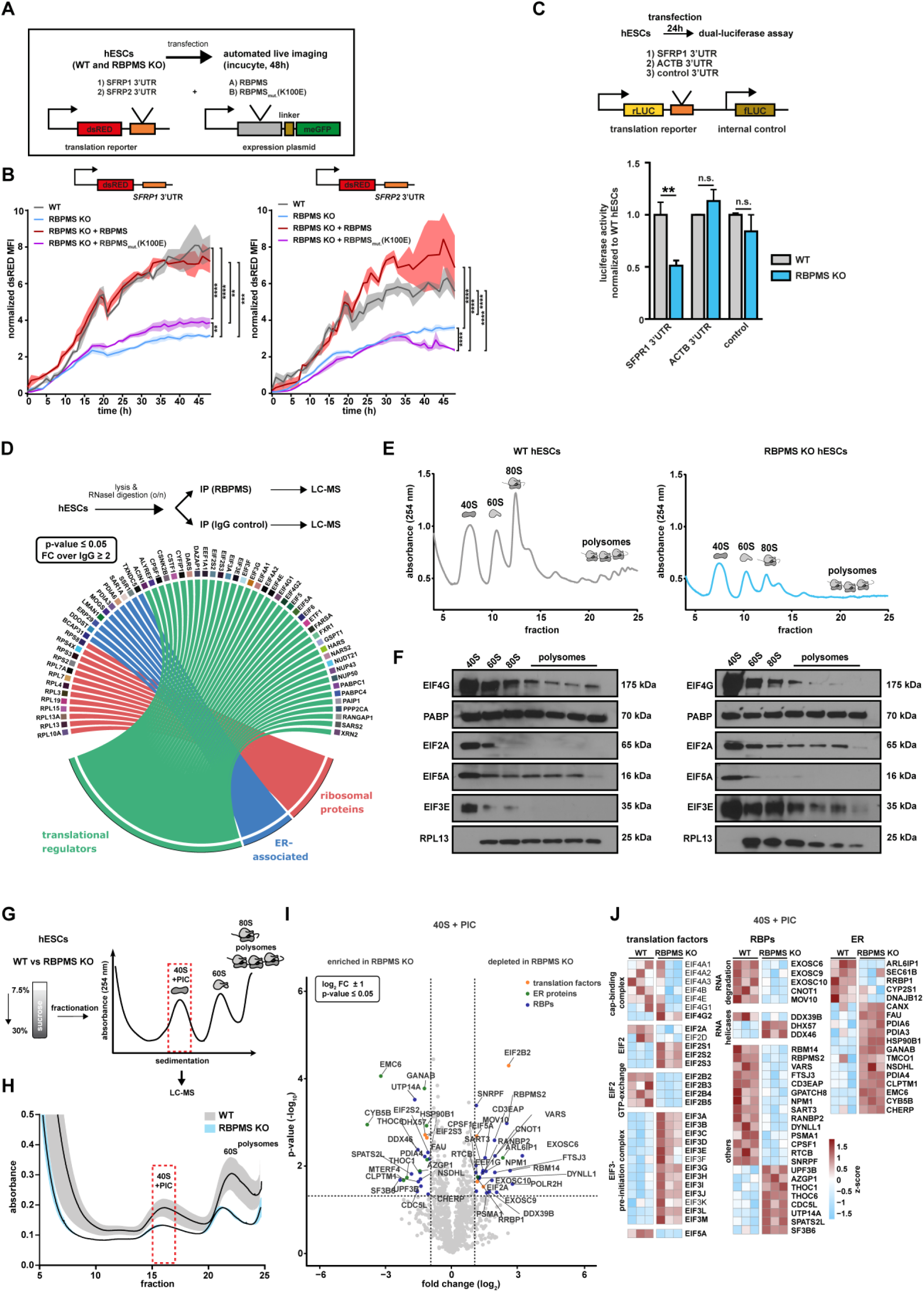
RBPMS specializes mRNA translation in pluripotency, selectively via 3’UTR binding and globally by shaping the composition of translation initiation complexes. **(A)** Schematic of the experimental workflow and destabilized dsRED-based reporter used to investigate the 3’UTR binding motif dependent regulation of translation by RBPMS in hESCs. Endogenous RBPMS binding motifs from *SFRP1* and *SFRP2* where used. **(B)** RBPMS activates translation of reporter mRNA carrying RBPMS binding motifs in the 3’UTR, evaluated by time-lapse microscopy. **(C)** Presence of RBPMS binding motif is required for 3’UTR binding dependent translation activation by RBPMS. Indicated luciferase based biscitronic reporters where transiently transfected in WT hESCs and evaluated as relative luciferase activity for the indicated constucts. Global impact of the loss of RBPMS on ribosome occupancy in hESCs, revealed by two-step regression analysis of the mRNAs enriching on ribosomal fractions. **(D)**, Schematic of the IP-MS workflow employed to characterize RBPMS complex. Circos plot (below) summarizes the key proteins specifically identified to immunoprecipitate with RBPMS and their role in translational control (immunoprecipitation of RBPMS after extended treatment with RNase I over IgG control followed by LC-MS) Log_2_ LFQ >25 in all three biological replicates and log_2_ fold change >S1, p-value ≤ 0.05). (n=3 unless otherwise indicated). Polysome profiling (E) followed by (F) western blot analysis to evaluate the distribution of the indicated translation factors in RBPMS-KO hESCs compared to WT. (n=2) **(G)** Schematic outlining the translation complex profiling based isolation of 40S and pre-initiation complex (PIC) followed by LC-MS in WT and RBPMS KO hESCs. **(H)** Translation complex profiling traces of WT and RBPMS KO hESCs (experiment performed in biological triplicates, shades represent SEM. Dashed red rectangle indicates fractions subjected to LC-MS/MS. **(I)** Proteins significantly changing in the 40S + PIC fraction between WT and RBPMS KO hESCs represented as Volcano plot. Dashed lines indicate significance thresholds (-log10 p-value ≥ 1.3 and log2 fold change ± 2 (selected translation factors, ER proteins and RBPs are highlighted by orange, green and blue dots, respectively). **(J)** Heatmap depicting differentially enriched translation factors, RBPs and translation-associated ER proteins in the 40S + PIC fraction between WT and RBPMS KO hESCs (significantly changing proteins are highlighted in bold) Error bars represent ±SEM; p-values calculated using Student’s t-test (*≤0.05, **≤0.01, ***≤0.001, ****≤0.0001, n = 3).

To obtain a further mechanistic understanding of RBPMS-mediated global translational control, we first performed immunoprecipitation of RBPMS from hESCs followed by mass spectrometry-based proteomics analysis after prolonged RNase I treatment (to avoid indirect RNA-mediated associations; **Figure 6D, Figure S9A-C** and **Table S6**). RBPMS co-purified with translational regulators, ribosomal proteins and proteins involved in ER-mediated translation including canonical regulators of translation initiation. Notably, EIF3 complex components, EIF5A and EIF4G, along with multiple ribosomal subunits, were specifically and significantly enriched with RBPMS, suggesting that RBPMS can influence the residence of key regulatory components on the translation apparatus of hESCs (**Figure 6D** and **Figure S9C).** This is in line with the enrichment of RBPMS on 40S complexes upon treatment with translation inhibitors and RNase (**Figure 1M**). Accordingly, we evaluated the distribution translation initiation factors interacting with RBPMS on ribosomal complexes upon RBPMS loss by polysome profiling followed by western blotting (**Figure 6E, 6F**). First, we specifically examined the enrichment of EIF4G (that mediates the crosstalk of the 43S preinitiation complex with EIF4F complexes) and of the poly(A)- binding protein PABP. Both displayed comparable levels indicating that RBPMS loss does not influence the predisposition of mRNAs to be translated (**Figure 6F**). However, a key component of the 43S preinitiation complex and a regulatory hub for global translation, EIF2A, showed aberrant retention across ribosomal fractions following RBPMS loss (**Figure 6F**) (Wek, 2018). Additionally, two key EIF3 complex components, EIF3E (involved in selective translation) and EIF3H (reported to be involved in selective translation during embryonic development) were aberrantly retained in polysomal fractions and markedly depleted from the 40S complex in RBPMS-KO cells, respectively (**Figure 6F**)(Choudhuri et al., 2013; Lin et al., 2020). Strikingly, EIF5A, essential for translation initiation, elongation, for error resolution at ribosomal pause sites, and for termination, was sequestered in the 40S ribosomal fraction in the absence of RBPMS (**Figure 6F**) (Schuller et al., 2017). The total levels of these factors do not reflect the change in their distribution pattern in ribosomal fractions in *RBPMS-*KO hESCs (**Figure S9D**). Encouraged by these data, next we asked whether the loss of RBPMS disrupts the abundance and assembly of translation initiation complexes on translationally engaged mRNAs in hESC. To this end, we sought to translation complex profiling (**Figure 6G, 6H**), a variant of polysome profiling which is specifically designed analyze translation initiation complexes (Wagner et al., 2020). To have an unbiased evaluation of the effect of the loss of RBPMS on translation initiation complexes we subjected the fraction containing 40S and preinitiation complex to LC-MS based proteomics analysis (**Figure 6I, Table S6)**. The loss of RBPMS disrupted the composition of translation initiation complexes including cap binding complex, EIF2 complex and EIF3 complex (**Figure 6J)**. Notably, components involved in GTP exchange in the EIF2 complex where significantly depleted while majority of the EIF3 complex where aberrantly retained on translation complexes upon RBPMS loss. As observed by the polysome fractionation based analysis (**Figure 6F**), EIF5A was significantly depleted (**Figure 6J)**. Notably, crucial mediators of ER-associated translation such as SEC61B and RRBP1 were significantly depleted due to the loss of RBPMS. In addition several RBPs, including RANBP2 which is known to enhance the translation of secretory proteins, and the components of RNA degradation machinery where depleted upon RBPMS loss while RNA helicases such as DHX57 and DDX46 where enriched in the absence of RBPMS. Thus, our data show loss of RBPMS causes translation initiation defects highlighted by aberrant retention of the EIF3 complex and depletion of EIF5A from mRNAs revealing its role in regulating global translation in hESCs.

Finally, to confirm that RBPMS levels determine the translation status in hESCs, its competence to mesoderm and cardiac commitment, and to rule out discrepancies stemming from genome engineering “off-target” effects in *RBPMS-*KO hESCs, we knocked-in an inducible copy of RBPMS using PiggyBac transposon-based genomic insertion (**Figure S9E, 9F**). Timely re-expression of the most prominent cytosolic isoform of RBPMS (the lowest band in RBPMS immunoblot; **Figure 1L**) in RBPMS*-*KO cells fully restored both ribosome occupancy defects **(Figure S9G**) and protein synthesis (**Figure S9H)**, including SFRP1 protein levels, an RBPMS 3’UTR target and WNT signaling mediator (**Figure S9I)**. Notably, RBPMS reexpression also restored mesoderm commitment capacity, now allowing *RBPMS-*KO cells to generate terminally differentiated cardiomyocytes (**Figure S9J**). In summary, our data confirm RBPMS as a functional RAP essential for global translation in hESCs and a selective translation activator of target mRNAs via 3’UTR binding.

## Discussion

The appropriate abundance and balance of cell fate-determining morphogen signaling components at the state of pluripotency is vital for the ability of hESCs to undergo accurate lineage decisions. This is especially important for mesoderm and cardiac mesoderm differentiation and patterning, where early exposure to WNT-BMP-NODAL signaling dosage encountered by the cells committing to mesoderm that transgresses the primitive streak is postulated to predetermine their ability to commit to future cardiac lineages both *in vitro* and *in vivo* (Ghimire et al., 2021; Jiang et al., 2022; Kempf et al., 2016; Loh et al., 2016; Rao et al., 2016). How this unique predisposition to a future terminal fate is molecularly regulated is currently unknown. Our work reveals that competence of hESCs for future cardiac commitment is predetermined already at the state of pluripotency in a specialized mRNA translational circuit controlled by RBPMS. Highlighting its key role in early embryonic cell fate decisions, RBPMS selectively primes the translation of regulatory components essential to initiate cardiac commitment program. Thus, it ensures the accurate abundance of ‘morphogen signaling infrastructure’ in hESCs that is essential to authorize the foundation of cardiac mesoderm upon receiving differentiation signals. Based on our findings, we propose that RBPMS is a translation specialization factor.

We postulate translation specialization as a regulatory mechanism that primes ribosomes to control translation temporally and/or spatially for a set of mRNAs necessary for future events in response to particular stimuli or fate transitions. This allows efficient division of labor among the ∼10M ribosomes present in each cell, which are tasked with synthesizing ∼2M proteins/min, so the flow of information is streamlined and, as we show, specialized.

Mechanistically, on the one hand, RBPMS associates with components of translation initiation complexes, and its loss abrogates translation initiation and ribosome recruitment, primarily by disrupting EIF2 and EIF3 complexes. Importantly, RBPMS loss severely depletes the translation apparatus of the key ‘surveillance’ factor EIF5A. These points to a role for RBPMS in shaping translation initiation in hESCs as its absence leads to global inhibition of translation. On the other hand, RBPMS selectively regulate translation of mesoderm instructive signal transduction components in hESCs by binding target 3’UTRs via its specific mRNA binding motif. In fact, our reporter assays showed that the insertion of RBPMS recognition elements in the 3’UTR suffices for boosting translation. Thus, reinstating RBPMS levels restores the mesoderm specification capacity of hESCs. Notably, not all RBPMS targets are translationally repressed. This is not uncommon for selective regulators of mRNA translation. For instance, the loss of eIF3D, a translation factor enabling selective translation upon stress, leads to both activation and inhibition of mRNAs in a context-dependent manner, in addition to its role as general translation initiation factor (Lee et al., 2015; Lee et al., 2016).

RNA binding proteins can affect distinct (or multiple) aspects of RNA processing including mRNA splicing, localization and translation in a cell type, developmental stage/context-dependent fashion (Hentze et al., 2018; Van Nostrand et al., 2020). For example, QKI is a regulator of mRNA splicing in cardiomyocytes while it is reported to regulate translation/ mRNA localization in astrocytes and germline lineages (Chen et al., 2021; Sakers et al., 2021). Thus, it is also possible that RBPMS controls distinct mRNA processing and regulatory mechanisms in cell type/ context- dependent fashion as reported in smooth muscle cells and recent mouse cardiomyocytes (Gan et al., 2022; Nakagaki-Silva et al., 2019). Importantly, the mRNA targets of RBPMS and its proposed role in pre-mRNA splicing in terminally differentiated mice cardiomyocytes remains to be experimentally determined. In contrast, in hESCs we reveal its role as a RAP selectively recruited in active ribosomes and controlling mRNA translation, without directly influencing transcription, mRNA splicing or mRNA abundance.

In summary, we propose a model by which the state of pluripotency is translationally poised for differentiation into future lineages via the specialized translation of the regulators of embryonic cell fate. Our work reveal that cell fate-specific translation specialization factors selectively program the translation of mRNAs encoding key developmental regulators that are essential to initiate future cell fate-choices, akin to how pioneering transcription factors program specific transcriptional networks allowing cell fate decisions. Collectively, we reveal a pivotal role for translational specialization in sculpting cellular identity during early developmental lineage decisions and propose that ribosomes act as a unifying hub for cellular decision-making rather than a constitutive protein synthesis factory.

## Material and Methods

### Human Pluripotent stem cells and culture conditions

HuES6 (genotype:female) cells (kindly shared by Boris Greber’s Lab, Max Planck Institute for Molecular Medicine (Muenster, Germany)), derived from inner cell mass of blastocysts, were used for the in vitro differentiation experiments. Cardiomyocyte reporter hiPSC line, WTC MYL7-GFP iPSC, was a gift from Mendjan lab, IMBA Vienna. Authenticated cell lines were provided by the indicated providers. hPSCs were maintained on Matrigel-coated 6-well dishes in FTDA medium (Frank et al., 2012). FTDA media contains DMEM/F-12 supplemented with 1X Penicillin/Streptomycin/Glutamine, 1X Insulin/Transferrin/Selenic Acid, 0.1% Human Serum Albumin, 1:100 lipid mix, 50 ng/ml FGF2, 0.2 ng/ml TGFβ1, 50 nM dorsomorphin and 4 ng/ml Activin A. For passaging, cells were washed with PBS and dissociated with Accutase, supplemented with 10 μM ROCK inhibitor (Y-27632) for 10 min at 37°C. Accutase was blocked with F12/DMEM and the desired number of cells were centrifuged for 2 min at 300 x g at RT. The cell pellet was resolved in 2 mL FTDA medium, supplemented with 10 μM Y-27632, and pipetted onto fresh Matrigel-coated plates followed by 24 h incubation at 37°C and 5% CO_2_. After 24 h, medium was changed to 2.5 mL fresh FTDA medium. Medium change was repeated daily with increasing FTDA volumes, until cells reached confluency.

## Methods Details

### RBPMS KO using CRISPR-Cas9 in human ESCs

For the generation of CRISPR-Cas9-mediated KO hPSCs, 175000 cells were seeded per well of a 6-well plate and incubated overnight. The following day, the transfection mix containing 200 µl OptiMEM, 12 µl FuGene HD and 3 µg px330A RBPMS gRNA plasmid was prepared. The transfection mix was vortexed for 15 sec and incubated 15 min at RT. While incubation, medium was replaced by 2 ml fresh FTDA growth medium. After 15 min, the transfection mix was added dropwise to the cells, followed by 24 h incubation. After 24h, GFP expression was assessed under the cell culture microscope. If the number of GFP-positive cells was sufficient (10% or higher), cells were selected for 24 h by adding 0.5 µg/ml puromycin to the FTDA growth medium. After 24 h, puromycin was removed from FTDA and cells were recovered for 2 to 3 days. For the selection of single-cell derived RBPMS KO clones, cells were seeded on clonal dilution (2000-6000 cells/well of a 6-well plate) in 1.5 FTDA growth medium with 10 µM Y-27632. After 24, 0.5 ml FTDA growth medium with 10 µM Y-27632 was added to the cells. The next day, 1ml of medium was replaced with 1ml fresh FTDA. From the following day, all medium was replaced daily by 1 to 1.5 ml FTDA growth medium until colonies were large enough for screening. For screening, 20% of each colony (preferably the middle part, splitting the colony in half) was scraped from the plate and lysed in 30µl of Quick extract DNA Extraction solution (Biozym, 101094). Cells were vortexed for 15 seconds and incubated for 6 min at 65°C, followed by an additional 15 sec vortexing and incubation at 98°C for 2 min. For screening PCRs (primers see Table 2), 5 µl of isolated genomic DNA (gDNA) were used, while the remaining gDNA was stored at -20°C. Positive clones were be grown for some more days until they reach a proper size for picking. For picking the colony was cleaned from surrounding cells and divided into small pieces by using a 27-G syringe. Colony pieces were transferred to a fresh well containing 2 ml FTDA growth medium with 10 µM Y-27632. After 1 week, colonies were divided into small pieces again and dissociated with Dispase. For this, cells were washed once with DPBS. After washing, 1 ml Dispase was added and cells were incubated for 5 min at 37°C. After 5 min incubation, colonies were carefully washed twice with DBPS. Then 2 ml FTDA growth medium was added and cells were scraped from the plate using a cell scraper. Colonies were centrifuged briefly to sediment at the bottom of the tube. The supernatant was removed and cells were seeded in onto a fresh 6-well plate. When the well was covered at least by 50% with colonies, cells were returned to monolayer culture by splitting with Accutase.

### Generation of stable inducible *RBPMS* expression lines

For the generation of doxycycline-inducible *RBPMS* expression in RBPMS KO hESCs, PiggyBac-based transposon-mediated genomic insertion was performed as described in Rao et al., 2016. Briefly, 175000 cells were seeded in Matrigel-coated 6-well dishes in FTDA medium supplemented with 10 μM Y-27632. The following day, a transfection mix (200 μl Optimem + 5 μl Lipofectamine3000© and the three plasmids containing the *RBPMS* construct, the transactivator and the transposase in a ratio of 10:3:1 with a total DNA amount of 2.8 μg) was prepared and incubated at 10s min at RT. Prior to adding the transfection mix, the medium was changed to fresh FTDA. After dropwise addition of the transfection mix, cells were incubated for 24h at 37°C. The next day, medium was changed to fresh FTDA. The following day cells were seeded in clonal density on 6-well plates (1000-2000 cells per well). From the day after seeding, medium was changed to fresh FTDA, supplemented with 0.5μg/ml G-418. Cells were selected for 10-14 days in FTDA and G-418 until single cell-derived colonies emerged. To test proper integration and expression of the integrated *RBPMS* construct, cells were induced by adding 20 ng/ml doxycycline to the growth medium and incubation for 24h at 37°C. The following day GFP positive cells were observed using a fluorescence microscope. For cardiac differentiation of the obtained clones, 50ng/ml doxycycline was added to the cardiac differentiation medium.

### Mesoderm differentiation protocol

For mesoderm differentiation, we adapted the protocol (Frank et al., 2012). Briefly, we seeded cells in FTA medium supplemented with 10 μM Y-27632 (see Maintenance of human pluripotent stem cells), which was lacking Dorsomorphin. Upon reaching 70% confluency, cells were treated with mesoderm induction medium (FTA + 5 µM CHIR and 5 ng/ml BMP4 for 24h at 37°C. Cells were collected for downstream experiments 24h after mesoderm induction.

### Induction of ectoderm and endoderm lineage differentiation

For neuroectoderm differentiation, cell aggregates were derived from confluent cell layers and cultured in suspension culture dishes in basic medium containing FGF2, Dorsomorphin and Y27632 (10 μM). Afterward, cells were treated for four days in neural induction medium (basic medium containing 0.5 μM PD0325901, 15 μM SB431542, 0.5 μM Dorsomorphin, and 10 μM Y27632). After four days, neural spheres were plated on Matrigel-coated plates and kept for another four days in the neural induction medium.

For induction of endoderm differentiation, 400000 cells were seeded in Matrigel-coated 6-well plates, in FTA supplemented with 10 μM Y-27632. The following day, cells were induced by adding endoderm induction medium (basic medium with 100 ng/ml Activin A, 10 ng/ml BMP4 and 5 μM CHIR99021) for 24h at 37°C. After 24h, medium was changed to basic medium containing 100 ng/ml Activin A. Cells were kept in this medium for an additional 72h.

### Cardiomyocyte differentiation

Cardiomyocyte differentiation was performed as previously described (Rao et al., 2016). Briefly, confluent hESCs were dissociated into single cells with Accutase at 37°C for 10 min. Accutase was stopped with two volumes of F12/DMEM. Cells were counted and 230000 cells per square cm (450000 for one well of a 24-well plate) were centrifuged for 2 min at 300 g at RT. The cell pellet was resolved in ITS medium, containing 1.5 μM CHIR and 1.5 ng/ml BMP4, and seeded on Matrigel-coated 24-well plates. ITS media comprised of Knockout DMEM, 1X Penicillin/Streptomycin/Glutamine,1X ITS supplement (1000x stock), 10 μM Y-27632, 25 ng/ml FGF2, 1-2 ng/ml BMP4 and 1-2 μM CHIR99021. To ensure equal distribution and attachment of cells, plates were moved crosswise, tapped several times and left outside for 20 min before transferring them to the incubator. After 24 h, medium was changed to TS medium. After 48 h, medium was changed to TS medium supplemented with 10 μM canonical Wnt-Inhibitor C59 for 48 h. TS media comprised of Knockout DMEM, 1X Penicillin/Streptomycin/Glutamine, 1X TS supplement, 250 μM 2-Phospho-L-Ascorbic Acid (TS supplement, 100X: 5.5 μg/ml Transferrin and 6.7 ng/ml Sodium Selenite in 100ml sterile PBS. After 48 h, medium was changed to fresh TS until beating cells were observed at around day 8. From then cardiomyocytes were cultivated Knockout DMEM supplemented with 2% FCS, L-Glutamine and Penicillin/Streptomycin until cells were used for downstream analysis.

### Cardioid differentiation

For the generation of cardiac organoids (cardioids), media and conditions were adapted from Hofbauer et al. (2021). Briefly, hPSCs grown in FTDA to approximately 70% confluency. For cardioid formation, 7500 cells/well were seeded into ultra-low-attachment 96-well plates (Corning) and centrifuged for 5 min at 200 g. After 24h cells were induced with FLyABCH medium (CDM containing 30 ng/ml FGF2 (Proteintech), 50 ng/ml Activin A (Miltenyi Biotech), 10 ng/ml BMP4 (R&D Systems) 3 µM CHIR (Tocris), 5 µm LY294002 (Tocris) and 1 µg/ml Insulin (Roche)) for 36-40h. After this, cells were treated with BFIIWPRa medium (CDM, containing 8 ng/ml FGF2, 10 ng/ml BMP4, 1 µM IWR-1 (Tocris), 0.5 µM retinoic acid (Sigma)) for 96h with media change every 24h. Following this, medium was changed to BFI (CDM, containing 10 ng/ml BMP4, 8 ng/ml FGF2 and 10 µg/ml Insulin) for 48h with media change after 24h. After 48h in BFI, cells were kept in CDM+I (10 µg/ml Insulin) until being harvested or used for imaging. For RNA isolation, 3 organoids were pooled per replicate and transferred into 250 µl Trizol solution. For imaging, organoids were fixed for 15 min in 4% PFA, washed three times in PBS and kept in at 4°C until further processing.

### OPP and puromycin labelling

For OPP labelling, the reagents were prepared according to the manufacturer’s instruction (ThermoFisher). Cells were seeded onto coverslips in FTDA. The following day cells were treated for 30 min at 37° C with 2 µM OPP added to the growth medium. After 30 min, cells were washed with PBS and fixed by 3.7% PFA for 15 min at RT. Following fixation, cells were permeabilized with 0.5% Triton for 15 min at RT. Afterwards cells were washed once with PBS and treated with OPP reaction cocktail for 30 min. After 30 min, cells were washed once with rinse buffer and nuclei were stained using NuclearMask™ Blue Stain for 30 min at RT in the dark. After washing, cells were ready for imaging analysis.

For puromycin labelling, cells were washed once with PBS and medium was changed to growth medium containing 0.5 µg/ml puromycin, while addition of 0.1 µg/ml cycloheximide was used as a negative control. After 15-30 min incubation at 37°C, cells were washed once with PBS and then harvested and flesh frozen for western blot or fixed with 4% PFA for immunostainings, respectively.

### Oxygen consumption rate (OCR) analysis by Seahorse

For OCR quantification, 35000 and 75000 hPSCs were seeded into Seahorse XFe96 (Seahorse Bioscience, Agilent) cell culture plates in FTDA growth medium and incubated overnight. One hour before assay, cells were treated with XF Base medium (Seahorse Bioscience, Agilent) supplemented with 1 mM sodium pyruvate, 2 mM L-glutamine, and 10 mM glucose (pH 7.4). The assay was set as follows: 3 measurements (3 min mixing plus 3 min measuring each) for basal respiration, 3 measurements after oligomycin (1 μM) injection, 3 measurements after FCCP (1 µM) injection and 3 measurements after rotenone/antimycin A (0.5 μM) injection.

### Cell cycle analysis by fluorescence-assisted cell sorting (FACS)

For fluorescence-assisted cell sorting (FACS), 1 M cells were harvested by Accutase and washed with PBS. After harvesting, cells were fixed in 1ml cold 70% EtOH by adding drop-wise to cell pellet and vortexing, followed by 30 min incubation on ice. After fixation, cells were centrifuged at 1000 x g 5 min at 4°C. The supernatant was aspired and cells were washed with twice with cold PBS. For RNA removal, 50 µl RNaseA solution were directly added to the pellet and cells were incubated for 30 min on ice. After RNase A treatment, 400 μl PI solution per million cells was directly added to cells in RNase A solution and mixed by pipetting. The mixture was incubated for 10 min at RT. Nine samples were analysed by flow cytometry in the Institute for Genetics FACS facility at the University of Cologne.

### Active Ribosome Complex Mass Spectrometry (ARC-MS)

For arc-ms the AHARIBO protein module (Immagina®) was adapted. Briefly, hESCs were grown in FTDA to 70% confluency, washed once with PBS and treated for 40 min with methionine-free growth medium (Thermo Scientific) containing 10% FBS, and 0.8 mM L-leucine to deplete methionine reserves. After 40 min, 10 µl AHA regent was added to the medium and cells were incubated at 37°C for 5 min, following an addition of 2.6 µl sBlock for 5 min at 37°C. Afterwards, cells were placed on ice and washed once with 1 ml cold PBS. PBS was removed with a pipette and cells were lysed using 40 µl cold lysis buffer by using a cell scraper. Cell lysate was transferred to a 1.5 ml microcentrifuge tube and cell debris was pelleted by centrifugation at 20,000 x g for 5 min at 4°C. The supernatant was transferred to a new tube and kept it on ice for 10 min. Absorbance was measured by Nanodrop at 260 nm with lysis buffer as blank subtraction. For the capture of active ribosome complexes, 0.4 AU were transferred to a new tube and the volume was adjusted to 100 µl with freshly prepared W buffer. To this, 100 µl dBeads were added and the mixture was incubated for 60 min at 4°C on a rotating wheel. After 60 min, the supernatant was removed and the beads were washed twice with UWS buffer. The supernatant of these washes (containing active ribosomes and associated proteins) was combined and stored at 4°C until mass spectrometry preparation via solution digest. For validation of AHA capture, two more washes in UWS were performed and beads were resuspended in 200 µl destilled water. For proteomics analysis, an on-bead digestion was performed.

### Polysome profiling and TCP isolation

Prior to lysis, cells were treated with FTDA growth medium containing 0.1 mg/ml cycloheximide for 5 min at 37°C and 5% CO2. After 5 min, medium was removed and cells were washed once with ice cold DPBS, containing 0.1 mg/ml cycloheximide. Then, cells were scraped from the dish and centrifuge at 300 x g for 2 min. The supernatant was removed and the pellet was subjected to lysis or stored at -80°C. For lysis, the cell pellet was resolved in 0.5 ml ice cold lysis buffer (50 mM HEPES pH 7.4, 100 mM KCl, 5 mM MgCl_2_, 0.1% Triton X,1 mM DTT, 100 µg/ml cycloheximide and EDTA-free protease inhibitor, SUPERaseIn™ RNase Inhibitor (adding 10 µl directly to lysate) and pipetted up and down several times. The lysate was transferred to 1.5 ml microfuge tubes and incubated on ice for 10 min. After 10 min, the solution was triturated 10 times through a sterile 27-G needle and clarified by centrifugation for 10 min at 20000 x g at 4°C. After centrifugation, the supernatant transferred to a new Eppendorf tube. For input RNA isolation 50 µl lysate were directly mixed with 500 ul of Trizol™ and stored at -80°C until RNA isolation. Prior to gradient preparation, the ultracentrifuge and rotor buckets were cooled down to 4°C. While cooling down, sucrose solutions (10 and 50% (w/v) sucrose, 100 mM HEPES pH 7.6, 100 mM KCl, 5 mM MgCl_2_, 1 mM DTT and 100 µg/ml cycloheximide in RNAse-free H_2_O) were prepared. Half of a SW-40 thin wall polypropylene centrifugation tube (SETON, #7031) was filled with 10% (w/v) sucrose solution (ca. 6 ml) by letting the solution slowly rinse down the wall of the tube. Then a syringe was filled with the 50% sucrose solution, which was slowly injected underneath the 10% sucrose solution. Excess 10% sucrose was removed and caps were added to the centrifugation tube. Gradient were generated with a Gradient Master™ (Biocomp, 100-003) at RT using the system pre- sets for continuous SW40Ti 10-50% sucrose gradients. After gradient preparation, caps were removed and the lysate was added on top of the gradients. The tubes were transferred to the rotor buckets and balanced to 0.01 g accuracy. After balancing, samples were centrifuged at 38000 rpm for 3 h at 4°C using low acceleration and deceleration settings in a SW40Ti rotor of a Beckman ultracentrifuge. For the fractionation, first the fractionator (Teledyne Isco) and fraction collector (Teledyne Isco) were assembled. 15 min prior to usage the UV detector was switched on. The tubings were first washed with ultrapure H2O and 70% ethanol to prevent bacterial contamination. Afterwards the system was equilibrated by pushing 70% sucrose solution (70% sucrose (w/v) in RNAse free water) until the injection needle. After equilibration, the centrifugation tube was attached to the fraction collector. The flow rate was set to 2 ml/min and fraction collection was initialized by starting the fractionation software. Collected fractions were transferred to ice and the profiles were analysed to mark the corresponding ribosomal fractions. Samples were either directly stored at -80°C for later protein isolation or mixed with 1 ml TRIzol™ reagent for RNA extraction and RNA sequencing.

In case of TS-Seq ERCC spike-in controls were added just before layering the lysate on the gradient. A second set of spike-in controls was added to the isolated by the sequencing core to assess the quantity of libraries.

For the isolation of the pre-initiation complex (TCP-MS) a protocol adapted from Herrmannová et al. (2020) was used. Briefly, per replicate 2x 10 cm dishes of 70% confluency were treated with cycloheximide (100 µg/ml final concentration) for 1 min at 37°C, washed with PBS and harvested by adding lysis buffer (10 mM HEPES [pH 7.5], 62.5 mM KCl, 2.5 mM MgCl_2_, 1 mM DTT, 1mM PMSF, 1 µg/ml Aprotinin, 1 µg/ml Leupeptin, 1 µg/ml Pepstatin, mini complete EDTA-free protase Inhibitor (Roche) 1% Triton X-100 and 100 µg/ml cycloheximide) by directly adding to plate. Lysates were incubated on ice for 5 min and cleared by centrifugation for 5 min at 16000 x g at 4°C. Supernatant was transferred to a new tube and absorbance was measured via Nanodrop. 12 AU were loaded on a 7-30% sucrose gradient (to the 30% sucrose solution 0,05% formaldehyde was added before gradient mixing). Lysates were either flash-frozen and storted at -80°C or separated by centrifugation for 14 h at 84000 x g at 4°C.

### Protein isolation from sucrose gradients fractions

For TS-MS and TCP MS proteins were isolated from by methanol chloroform precipitation (Friedman, D.B. and Lilley K.S. “Quantitative proteomics for two-dimensional gels using difference gel electrophoresis (DIGE) technology” in John M Walker, Protein Protocols, 3rd Edition, Humana Press, 2009.). Briefly, to 500 µl fraction 3 volumes (1500 µl) of water, 4 volumes methanol (2000 µl) and 1 volume (500 µl) chloroform were added, followed by vigorous vortexing and centrifugation at 14000 x g for 10 min at 4°C. After centrifugation, the upper layer, containing methanol and H_2_O was removed and 2000 µl fresh methanol was added. The solution was vortexed and centrifuged again. After removing the supernatant, the pellet was dried briefly. For proteomics sample preparation, the pellet was directly reconstituted in 8M Urea buffer, while for western blot analysis the pellet was reconstituted in 1x SDS loading buffer and denatured for 5 min at 95°C prior to loading onto the SDS gel.

### RNA Isolation and DNaseI treatment

RNA extraction was performed by using phenol-based TRIzol™ reagent (Thermo Fisher Scientific, 15596026) according to the manufacturer’s protocol. At the end of RNA extraction the RNA pellet was air-dryed at least 10 min at RT. The RNA was resolved in 10-30 µl (depending on the pellet size) RNase-free water. For the estimation of RNA concentration and purity, 1.0 µl RNA was measured via Nanodrop. After measurement RNA was either subjected to downstream experiments or stored at -80°C. For the removal of genomic DNA contamination, 15 µl RNA were mixed with 3 µl genomic DNase (Fermentas, EN0521), 3 10x DNase reaction buffer (Fermentas, EN0521) and filled to a total of 30 µl with RNAse free water. The reaction mix was incubated for 30 min at 37°C on a PCR cycler. After 30 min, 3µL of 50mM EDTA were added followed by 10 min incubation at 65°C. Following DNase treatment, the RNA was transferred back to ice and the concentration was measured again. Purified RNA was either directly reverse-transcribed or stored at -80°C.

### Enhanced cross-linking immunoprecipitation followed by sequencing (eCLIP-seq)

Enhanced cross-linking with immunoprecipitation was performed as described in Van Nostrand et al., 2016. Briefly, hPSCs (20 million cells) were UV-crosslinked (400 mJ/cm2 constant energy), lysed in iCLIP lysis buffer and sonicated (BioRuptor). Lysates were treated with RNase I (Ambion, AM2294) to fragment RNA, after which RBPMS protein-RNA complexes were immuno precipitated using the indicated antibody. In addition to the RBP-IPs a parallel size-matched input (SMI) library was generated for each sample; these samples were not immunoprecipitated with anti-RBPMS antibodies, but were otherwise treated identically. Stringent washes were performed as described in iCLIP, during which RNA was dephosphorylated with FastAP enzyme (Fermentas) and T4 PNK (NEB, M0201S). Subsequently, a 3′ RNA adaptor was ligated onto the RNA with T4 RNA ligase (NEB, M0242S). Protein-RNA complexes were run on an SDS-PAGE gel, transferred to nitrocellulose membranes, and RNA was isolated off the membrane identically to standard iCLIP. After precipitation, RNA was reverse transcribed with AffinityScript reverse transcriptase (Agilent, 600107), free primer was removed with ExoSap-IT (Thermo Fisher Scientific, 78201.1.ML), and a 3′ DNA adaptor was ligated onto the cDNA product with T4 RNA ligase (NEB). Libraries were then amplified with 2x Q5 PCR mix (NEB).

Purified libraries were then sequenced via HiSeq 3000 (Illumina) with 75 bp paired-end reads at the Cologne Center for Genomics (CCG).

### Microscopy Imaging

Cells were seeded on Matrigel or gelatine-coated coverslips or chamberslides (Ibidi) in respective growth medium. For fixation cells washed one time with PBS and fixed with 4% PFA for 10 min at RT. Post-fixation, the PFA was removed and the cells were washed three times with PBS. Cells in each chamber were treated for 10 mins with 1% Triton X-100 (in PBS) for permeabilization. Cells were then incubated with the blocking solution containing 2% BSA and 2% glycine in PBS- T or TBS-T for phosphor antibodies. After blocking, the blocking solution was removed and cells were washed one time with PBS-T/TBS-T. Cells were incubated with either single or double primary antibodies (different host species) in 0.5% BSA in PBST overnight at 4°C. Primary and secondary antibodies were incubated for 1 hour at RT or overnight at 4°C. After incubation in secondary antibody, cells were washed with PBS, while DAPI was added to the second wash. Antibodies used were listed in Key Resources Table. After washing samples were rinsed with water and mounted with ProLong Gold mounting solution. Images were acquired with Leica SP7 or SP8 confocal microscopes with 3x line averaging bi-directional scanning using a 63x oil objectives.

For live cell imaging, 50000 cells per well were seeded in 24 well dish coated with matrigel. The florescence readout was measured in IncuCyte S3. 16 areas in each well were imaged every 1 hour over a period of 48 hours after transfection.

### RNA sequencing and analysis

RNA was sequenced in the Cologne Center for Genomics and prepared according to the Illumina RNA Sequencing library preparation kit protocol. Libraries were sequenced on a HiSeq 3000 or NovaSeq Sequencers (Illumina), with stranded paired-end reads of 75bp read length with a depth of at least 5 M reads for TS-seq or 20 M reads for total RNA sequencing. Duplicated reads and short reads were discarded.

Differentially expressed analyses was performed on the RNA-Seq data during cardiac differentiation, including total RNA-seq in human, polyA RNA-seq in human and total RNA-seq in mouse. Reads were aligned to the hg19 genome using Star with the Gencode annotaion as the reference transcriptome. Sequences aligned to tRNA and rRNA genes were removed. Differential expression analyses and gene quantification was performed with Deseq2 Differential expression was identified using Deseq2 8(FDR< 0.01 and FC > 2). For analysis, genes filtered by Deseq automatic independent filtering for low normalized read counts were discarded from analysis.

Significant differential alternative splicing was identified with rMATS (version 4.1.2 doi: 10.1073/pnas.1419161111). Analysis of inclusion level across timepoints was performed by re- calculating percent inclusion (PSI) using junction-spanning reads only, requiring the read extend at least 10nt into the exon regions on both sides of the junction. Unless otherwise noted, at least 20 junction-spanning reads were required for calculating PSI values, and only events meeting this criteria in at least 2 replicates were included in further analysis.

### Proteomics sample preparation and LC/MS conditions

For in solution digest, the cell pellet was lysed in 8M Urea buffer. If proteins were already in solution (TS-MS or ARC-MS) an appropriate amount of Urea buffer was added to reach ≥ 6 M Urea. For whole cell proteomes, chromatin was degraded using a Bioruptor (5 min, cycle 30/30 sec). Lysates were centrifuged 15 min at 20.000g, and supernatant was transferred to a new tube. Protein concentration was measured using Bradford and at least 10 μg protein was used for further sample preparation. Dithiothreitol (DTT, 500 mM stock) was added to a final concentration of 5 mM and samples were incubated at 37°C for 1 h. After 1h, Chloroacetamide (IAA, 500 mM stock) to a final concentration of 40 mM was added, samples were vortexed and incubated in the dark for 30 min. Endoproteinase Lys-C was added at an enzyme:substrate ratio of 1:100 and samples were incubated at 37°C for 4 h. After 4h, samples were diluted with 50 mM TEAB to reduce urea concentration to 2 M. Trypsin was added at an enzyme:substrate ratio of 1:100 and samples were incubated at 37°C overnight. The following day, digestion was stopped by acidifying with formic acid with a final concentration of 1%. Samples were now ready for StageTip purification. For StageTip conditioning, two layers of C18 were stacked into a 0.2 ml pipette tip (=StageTip). For washing, 20 μl methanol were added to the StageTip, followed by centrifugation at 2.600 rpm for 2 min. Afterwards, 20 μl buffer B was added to the StageTip, followed by centrifugation at 2.600 rpm for 2 min. Following this, StageTip was washed twice with buffer A with a centrifugation at 2.600 rpm for 2 min. After the second wash, approximately 4 μl of buffer A were kept on top of the C18 material. Samples (acidified with formic acid) were loaded onto the StageTips, followed by centrifugation at 2.600 rpm for 4 min. This was repeated until all samples was loaded onto the StageTip. After loading, StageTips were washed with 20 μl buffer A and centrifuged at 2.600 rpm for 2 min. After washing, the StageTips were dried with a syringe and stored at 4°C until mass spectrometry runs were performed.

In case of on bead digest (ARC-MS) UREA was added to the beads in 100 μl destilled H_2_O to a concentration of 6 M. All subsequent steps were performed as described for the in solution digest. At the last step, tubes were placed on a magnetic rack and supernatant carefully transferred to a new tube.

In case of in gel digest, gel pieces were washed twice with 100 µl 50 mM ABC/50% Acetonitril. Afterwards, gel slices were dehydrated by incubating for 10 min in 100 μl Acetonitril. The supernatant was discarded and gel pieces were dried in a SpeedVac for 5 min at RT. The supernatant was discarded and gel slices were washed twice with 300μl of 20mM NH_4_HCO_3_ for 15 min at RT, followed by a wash with 300μl of 20 mM NH_4_HCO_3_ / CH_3_CN (50:50 v/v) for 15 minutes. Following washing, 100μl of CH_3_CN were added to dehydrate the gel pieces for 5 minutes. The supernatant was discarded and gel slices were dried in a SpeedVac. For reduction of proteins, gel pieces were incubated for 45 min in 100 µl 10 mM DTT at 56°C in a thermomixer. The supernatant was discarded, and the gel pieces were dehydrated by incubating in 100 µl acetonitrile for 10 min at RT. After discarding the supernatant, samples were alkylated by incubating in 100 µl 55 mM chloracetamide for 30 min at RT in darkness. After alkylation, gel slices were washed for 15 min with 100 μl 50 mM ABC and dehydrated by incubating in 100 μl Acetonitril for 15 min. Afterwards, gel pieces were dried with the SpeedVac for 5 min. For digestion, 3 ng/ µl (90% trypsin/10% LysC) was added on the gel pieces and they were incubated for 30 min at 4°C. Excess solution was removed, and 50 mM ABC was added to cover gel pieces, followed by digestion overnight at 37°C. The next day, supernatant was transferred to a new reagent tube. Peptides were extracted from the gel by adding 100 µl 30% acetonitrile/ 3 % trifluoroacetic acid for 20 min at RT. The solution was combined with the supernatant from before and 100 µl 100% acetonitrile was added to the gel pieces. After 20 min, the solution was removed and added to the solution from the previous steps. The solution was dried down in the SpeedVac to a volume ≤ 50 µl at a temperature ≤ 30°C. The sample was acidified by adding formic acid to a final concentration of 1% and was now ready for loading onto the StageTip as described above.

All samples were measured in the CECAD proteomics facility. For total proteomes, ARC-MS and TS-MS, samples were analyzed on a Q Exactive Plus Orbitrap mass spectrometer that was coupled to an EASY nLC (both Thermo Scientific). Peptides were loaded with solvent A (0.1% formic acid in water) onto an in-house packed analytical column (50 cm length, 75 µm I.D., filled with 2.7 µm Poroshell EC120 C18, Agilent). Peptides were chromatographically separated at a constant flow rate of 250 nL/min using the following gradient: 7-23% solvent B (0.1% formic acid in 80 % acetonitrile) within 35.0 min, 23-32% solvent B within 5.0 min, 32-85% solvent B within 5.0 min, followed by washing and column equilibration. The mass spectrometer was operated in data- dependent acquisition mode. The MS1 survey scan was acquired from 300-1750 m/z at a resolution of 70,000. The top 10 most abundant peptides were isolated within a 1.8 Th window and subjected to HCD fragmentation at a normalized collision energy of 27%. The AGC target was set to 5e5 charges, allowing a maximum injection time of 108 ms. Product ions were detected in the Orbitrap at a resolution of 35,000. Precursors were dynamically excluded for 20.0 s.

For IP-MS, all samples were analysed on a Q Exactive Plus Orbitrap (Thermo Scientific) mass spectrometer that was coupled to an EASY nLC (Thermo Scientific). Peptides were loaded with solvent A (0.1% formic acid in water) onto an in-house packed analytical column (50 cm length, 75 µm I.D., filled with 2.7 µm Poroshell EC120 C18, Agilent). Peptides were chromatographically separated at a constant flow rate of 250 nL/min using the following gradient: 4-6% solvent B (0.1% formic acid in 80 % acetonitrile) within 1.0 min, 6-30% solvent B within 200.0 min, 30-50% solvent B within 28.0 min, 50-95% solvent B within 1.0 min, followed by washing and column equilibration. The mass spectrometer was operated in data-dependent acquisition mode. The MS1 survey scan was acquired from 300-1750 m/z at a resolution of 70,000. The top 10 most abundant peptides were isolated within a 1.8 Th window and subjected to HCD fragmentation at a normalized collision energy of 27%. The AGC target was set to 5e5 charges, allowing a maximum injection time of 55 ms. Product ions were detected in the Orbitrap at a resolution of 17,500. Precursors were dynamically excluded for 40.0 s.

Mass spectrometric raw data were processed with Maxquant (version 1.5.3.8) using default parameters. Briefly, MS2 spectra were searched against the canonical Uniprot Human.fasta (downloaded at: 16.06.2017) database, including a list of common contaminants.

False discovery rates on protein and PSM level were estimated by the target-decoy approach to 1% (Protein FDR) and 1% (PSM FDR) respectively. The minimal peptide length was set to 7 amino acids and carbamidomethylation at cysteine residues was considered as a fixed modification. Oxidation (M) and Acetyl (Protein N-term) were included as variable modifications. The match-between runs option was enabled. LFQ quantification was enabled using default settings.

### Quantification and Statistical Analysis

All data presented here is from at least three independent experiments. Graphs were plotted using GraphPad Prism and R Stats packages. The Student’s t-test was used to test for significance. Mean values ± SEM are shown. Differences between more than 2 groups were tested by one-way ANOVA analysis. Symbols representing p value cut-offs in the figures i.e., ^∗^,^∗∗^ and ^∗∗∗^ refers to p values ≤ 0.05, ≤ 0.01 and ≤ 0.001 respectively.

## RESOURCES TABLE

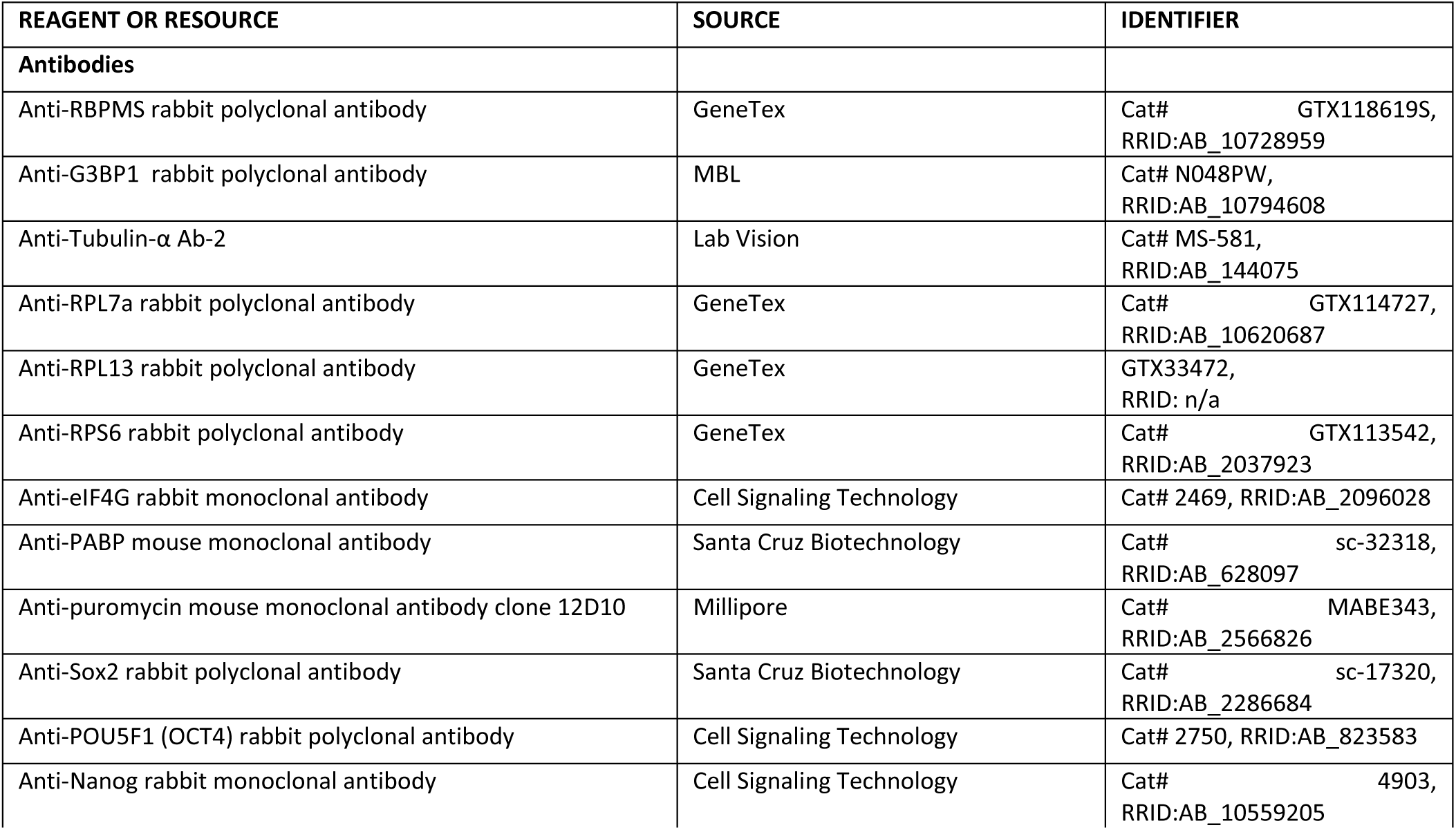
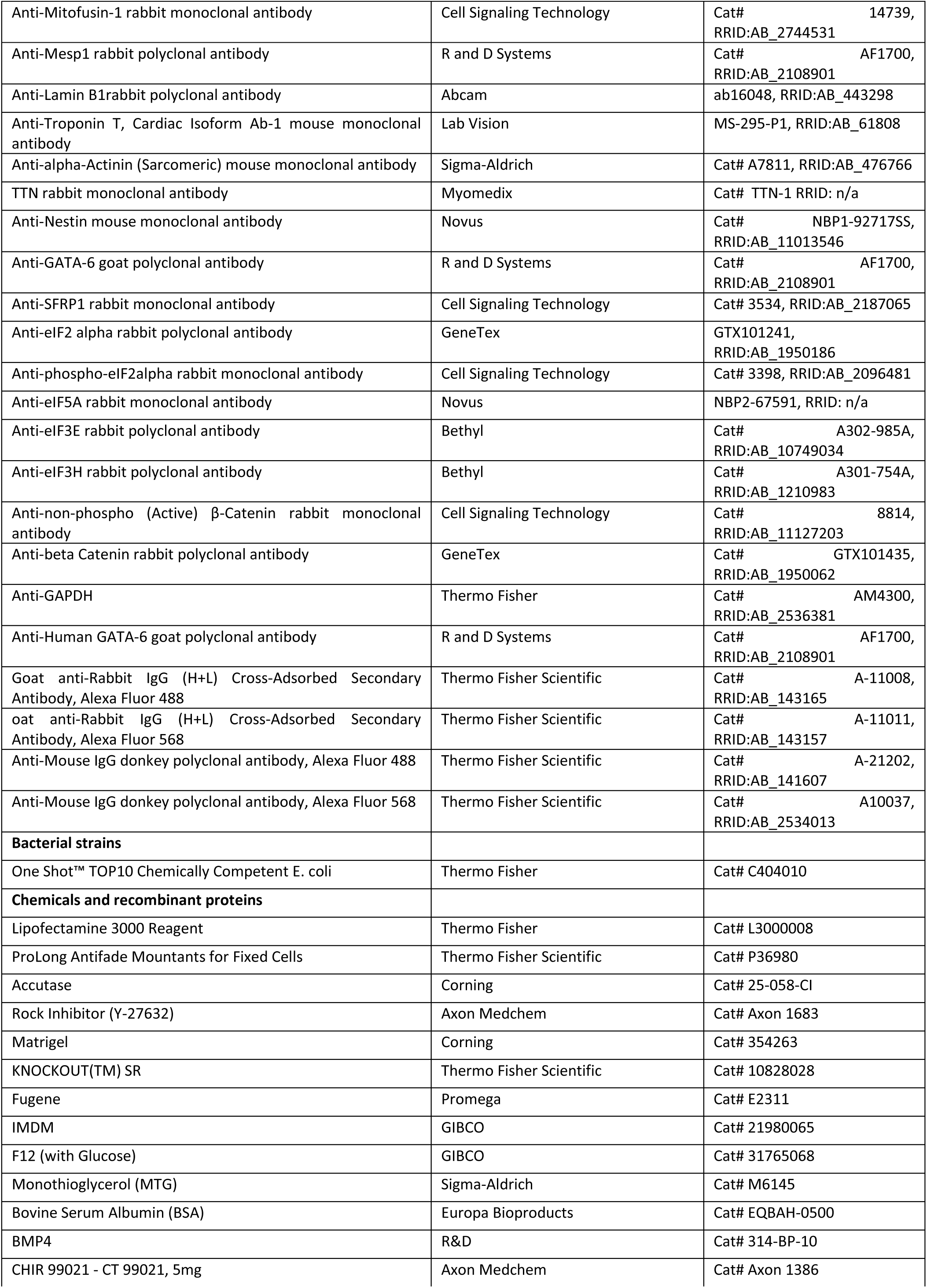
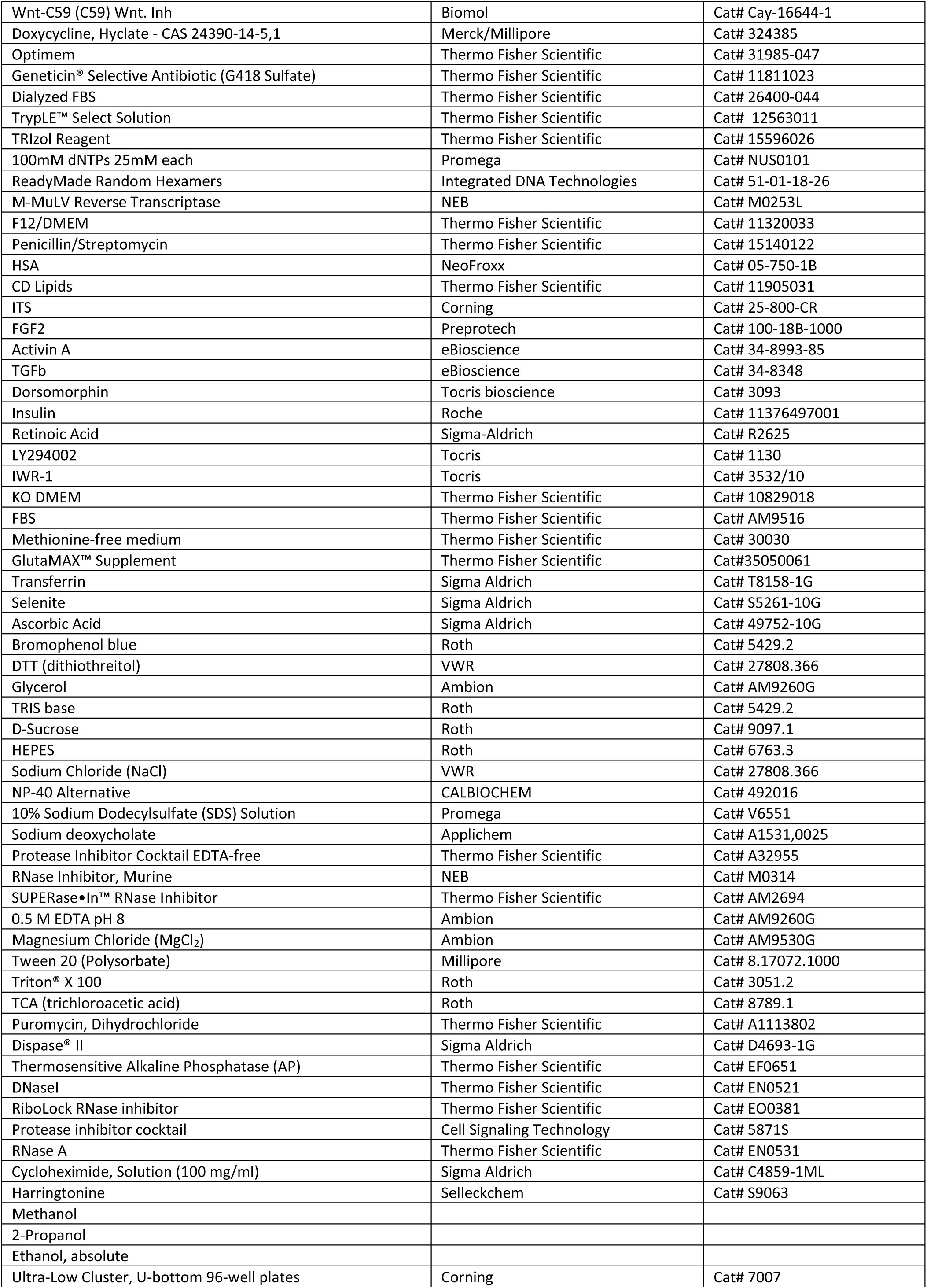
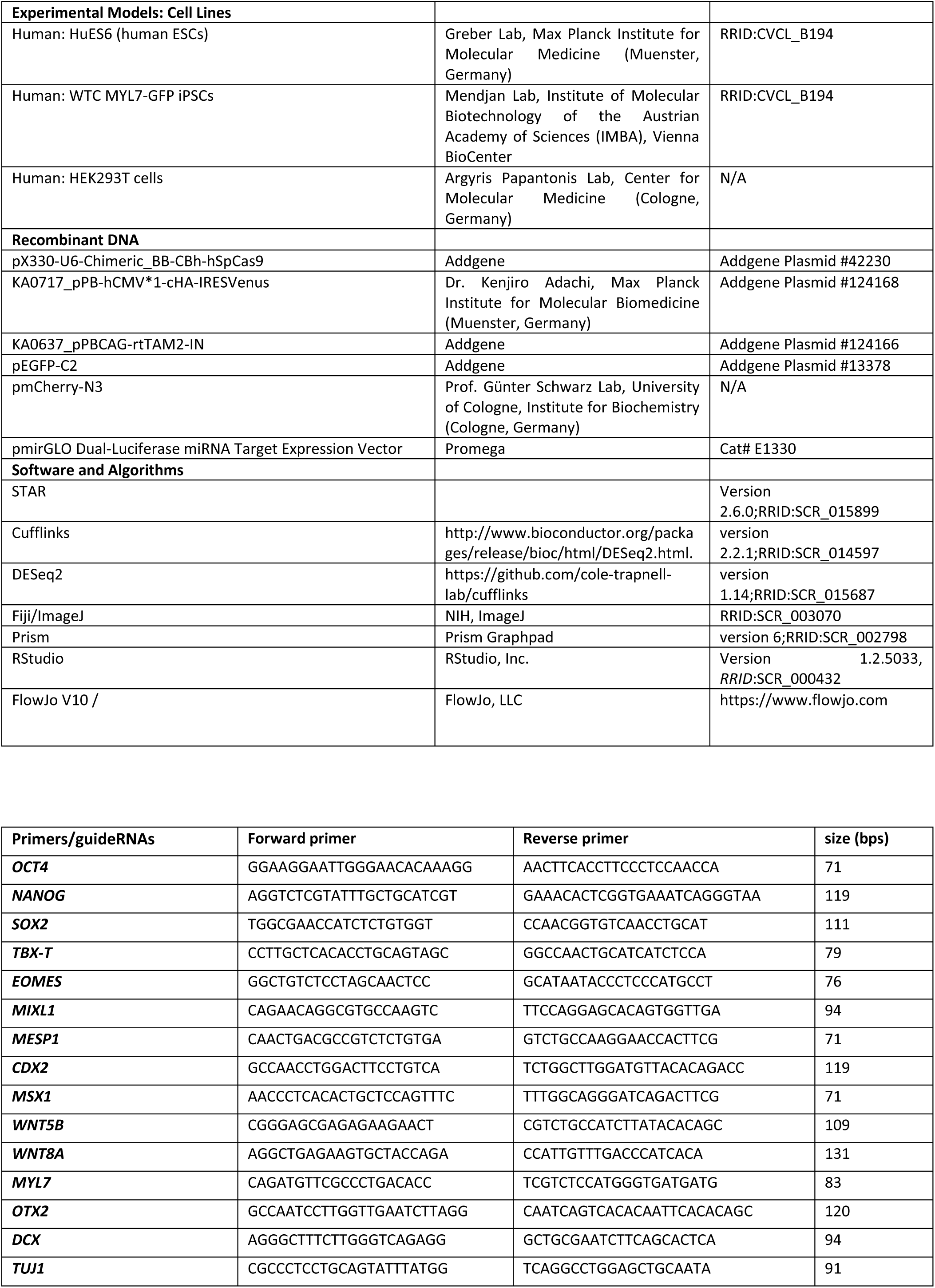
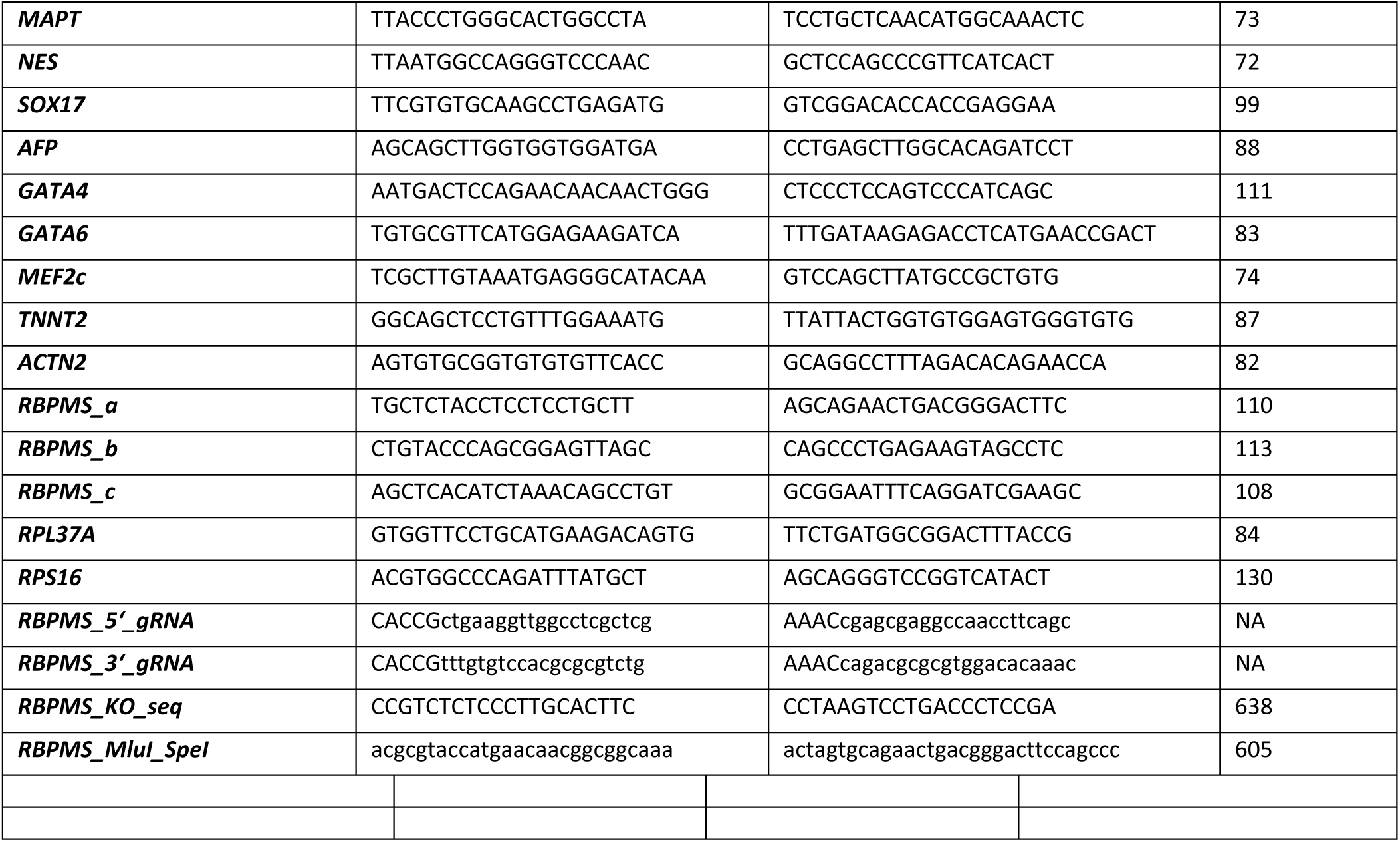

### DNA sequences

#### *SFRP1* binding site

ctcgagGGCAGGGTGGGGAGGGAGCCTCGGGTGGGGTGGGAGCGGGGGGGACAGTGCCCCGGG AACCCGGTGGGTCACACACACGCACTGCGCCTGTCAGTAGTGGACATTGTAATCCAGTCGGC TTGTTCTTGCAGCATTCCCGCTCCCTTCCCTCCATAGCCACGCTCCAAACCCCAGGGTAGCCA TGGCCGGGTAAAGCAAGGGCCATTTAGATTAGGAAGGTTTTTAAGATCCGCAATGTGGAGCA GCAGCCACTGCACAGGAGGAGGTGACAAACCATTTCCAACAGCAACACAGCCACTAAAACA CAAAAAGGGGGATTGGGCGGAAAGTGAGAGCCAGCAGCAAAAACTACATTTTGCAACTTGTT GGTGTGGATCTATTGGCTGATCTATGCCTTTCAACTAGAAAAtctagatctagaa

#### *SFRP2* binding site

TCCCGGCATCCTGATGGCTCCGACAGGCCTGCTCCAGAGCACGGCTGACCATTTCTGCTCCGG GATCTCAGCTCCCGTTCCCCAAGCACACTCCTAGCTGCTCCAGTCTCAGCCTGGGCAGCTTCC CCCTGCCTTTTGCACGTTTGCATCCCCAGCATTTCCTGAGTTATAAGGCCACAGGAGTGGATA GCTGTTTTCACCTAAAGGAAAAGCCCACCCGAATCTTGTAGAAATATTCAAACTAATAAAAT CATGAATATTTTTATGAAGTTTAAAAATAGCTCACTTTAAAGCTAGTTTTGAATAGGTGCAAC TGTGACTTGGGTCTGGTTGGTTGTTGTTTGTTGTTTTGAGTCAGCTGATTTTCACTTCCCACTG AGGTTGTCATAACATGCAAATTGCTTCAATTTTCTCTGTGGCCCAAACTTGTGGGTCACAAAC CCTGTTGAGATAAAGCTGGCTGTTATCTCAACATCTTCATCAGCTCCAGACTGAGACTCAGTG TCTAAGTCTTACAACAATTCATCATTTTATACCTTCAATGGGAACTTAAACTGTTACATGTATC ACATTCCAGCTACAATACTTCCATTTATTAGAAGCACATTAACCATTTCTATAGCATGATTTCT TCAAGTAAAAGGCAAAAGATATAAATTTTATAATTGACTTGAGTACTTTAAGCCTTGTTTAAA ACATTTCTTACTTAACTTTTGCAAATTAAACCCATTGTAGCTTACCTGTAATATACATAGTAGT TTACCTTTAAAAGTTGTAAAAATATTGCTTTAACCAACACTGTAAATATTTCAGATAAACATT ATATTCTTGTATATAAACTTTACATCCTGTTTTACCTA

#### *ACTbeta* (chicken) 3’UTR

ctcgagAGACCGGACTGTTACCAACACCCACACCCCTGTGATGAAACAAAACCCATAAATGCGCATAAAACAAGACGAGATTGGCATGGCTTTATTTGTTTTTTCTTTTGGCGCTTGACTCAGGATTAAAAAACTGGAATGGTGAAGGTGTCAGCAGCAGTCTTAAAATGAAACATGTTGGAGCGAACGCCCCCAAAGTTCTACAATGCATCTGAGGACTTTGATTGTACATTTGTTTCTTTTTTAATAGTCATTCCAAATATTGTTATAATGCATTGTTACAGGAAGTTACTCGCCTCTGTGAAGGCAACAGCCCAGCTGGGAGGAGCCGGTACCAATTACTGGTGTTAGATGATAATTGCTTGTCTGTAAATTATGTAACCCAACAAGTGTCTTTTTGTATCTTCCGCCTTAAAAACAAAACACACTTGATCCTTTTTGGTTTGTCAAGCAAGCGGGCTGTGTTCCCCAGTGATAGATGTGAATGAAGGCTTTACAGTCCCCCACAGTCTAGGAGTAAAGTGCCAGTATGTGGGGGAGGGAGGGGgtcgac

#### RBPMS CDS

atgaacaacggcggcaaagcCGAGAAGGAGAACACCCCGAGCGAGGCCAACCTTCAGGAGGAGGAGGTCCGGACCCTATTTGTCAGTGGCCTTCCTCTGGATATCAAACCTCGGGAGCTCTATCTGCTTTTCAGACCATTTAAGGGCTATGAGGGTTCTCTTATAAAGCTCACATCTAAACAGCCTGTAGGTTTTGTCAGTTTTGACAGTCGCTCAGAAGCAGAGGCTGCAAAGAATGCTTTGAATGGCATCCGCTTCGATCCTGAAATTCCGCAAACACTACGACTAGAGTTTGCTAAGGCAAACACGAAGATGGCCAAGAACAAACTCGTAGGGACTCCAAACCCCAGTACTCCTCTGCCCAACACTGTACCTCAGTTCATTGCCAGAGAGCCATATGAGCTCACAGTGCCTGCACTTTACCCCAGTAGCCCTGAAGTGTGGGCCCCGTACCCTCTGTACCCAGCGGAGTTAGCGCCTGCTCTACCTCCTCCTGCTTTCACCTATCCCGCTTCACTGCATGCCCAGATGCGCTGGCTCCCTCCCTCCGAGGCTACTTCTCAGGGCTggaagtcccgtcagttctgc

#### RBPMSmut CDS

atgaacaacggcggcaaagcCGAGAAGGAGAACACCCCGAGCGAGGCCAACCTTCAGGAGGAGGAGGTCCGGACCCTATTTGTCAGTGGCCTTCCTCTGGATATCAAACCTCGGGAGCTCTATCTGCTTTTCAGACCATTTAAGGGCTATGAGGGTTCTCTTATAAAGCTCACATCTAAACAGCCTGTAGGTTTTGTCAGTTTTGACAGTCGCTCAGAAGCAGAGGCTGCAAAGAATGCTTTGAATGGCATCCGCTTCGATCCTGAAATTCCGCAAACACTACGACTAGAGTTTGCTGAGGCAAACACGAAGATGGCCAAGAACAAACTCGTAGGGACTCCAAACCCCAGTACTCCTCTGCCCAACACTGTACCTCAGTTCATTGCCAGAGAGCCATATGAGCTCACAGTGCCTGCACTTTACCCCAGTAGCCCTGAAGTGTGGGCCCCGTACCCTCTGTACCCAGCGGAGTTAGCGCCTGCTCTACCTCCTCCTGCTTTCACCTATC CCGCTTCACTGCATGCCCAGATGCGCTGGCTCCCTCCCTCCGAGGCTACTTCTCAGGGCTggaagtcccgtcagttctg

## Supporting information

Bartsch et al 2022 Supplementary files

## Acknowledgments

We acknowledge the central facilities at CECAD and CMMC for technical assistance and Annibaldi lab and Lucardi lab for advice in various stages of this project. We thank Prof. Gene Yeo (UCSD) and Asst. Prof Eric Van Nostrand (Baylor College) for sharing the eCLIP technology, Van Nostrand lab for the mRNA splicing analysis and Prof. Leoš Shivaya Valášek (The Czech Academy of Sciences) for sharing the TCP-profiling method. We would like to thank Dr. Janine Altmueller, CCG-Cologne and Immagina Biotech for advice and assistance.

## Author contributions

Conceptualization: LK

Methodology: DB, GA, KK, MC, SM, LK

Investigation: DB, LK

Visualization: DB

Supervision: LK

Writing—original draft: LK

Writing—review & editing: LK, DB, HB, AP

## Data and materials availability

Data from all proteomics experiments including ARC- MS and TCP-MS are available via ProteomeXchange with identifier PXD032045. Time course transcriptomics data set, related to Figure 3, is available in GEO: GSE205300, eCLIP-seqdata in GSE205794 and TS-seq data in GSE205793. “

