## Supplementary material for "A specialized mRNA translation circuit instated in pluripotency presets the competence for cardiogenesis in humans": Bartsch et al 2022 Supplementary files

**This PDF file includes:**

Figs. S1 to S9

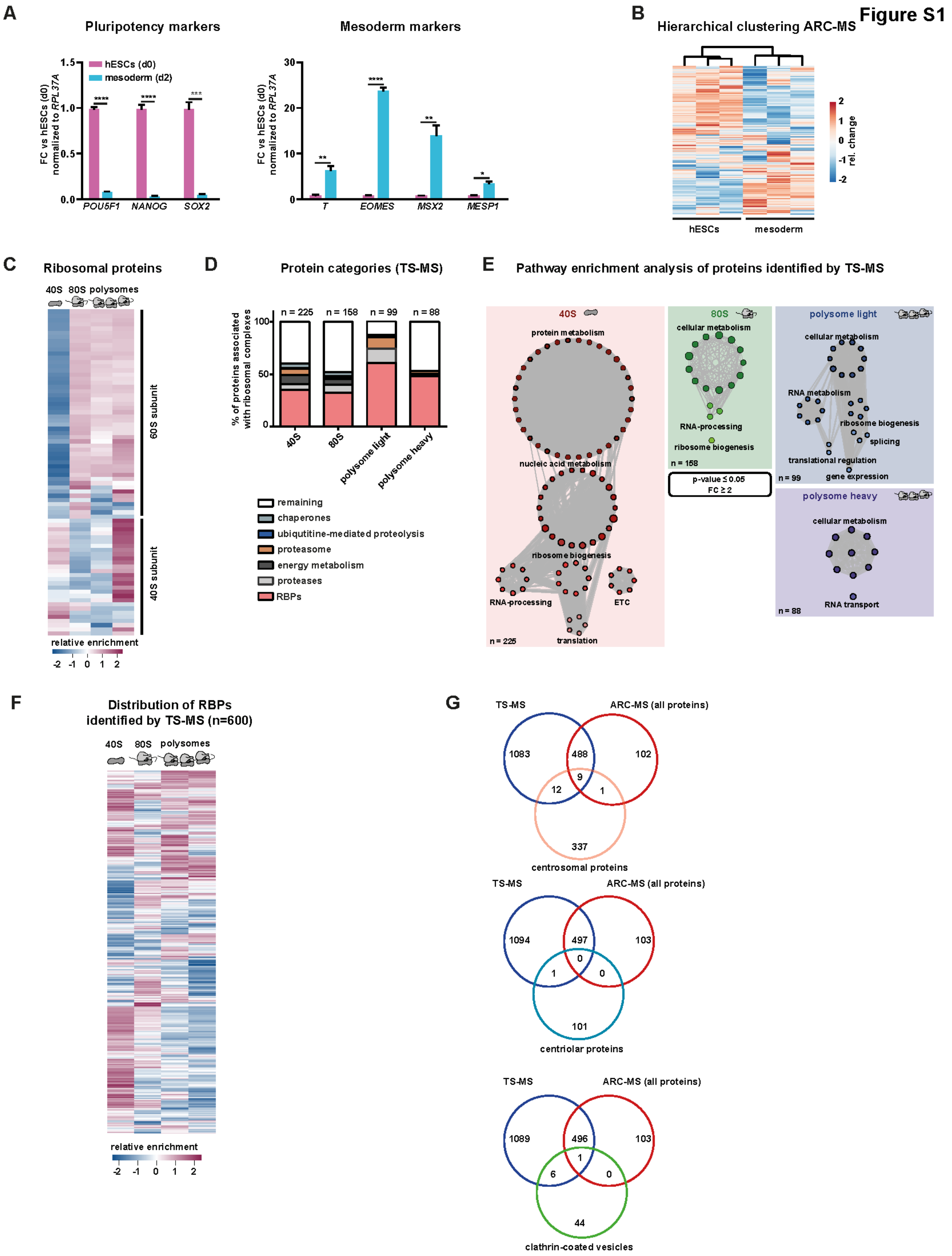
Figure S1
A
Pluripotency markers
hESCs (d0)
mesoderm (d2)
FC vs hESCs (d0) normalized to RPL37A
POU5F1
NANOG
SOX2
Mesoderm markers
T
EOMES
MSX2
MESP1
B
Hierarchical clustering ARC-MS
rel. change
hESCs
mesoderm
C
Ribosomal proteins
40S 80S polysomes
60S subunit
40S subunit
relative enrichment
D
Protein categories (TS-MS)
% of proteins associated with ribosomal complexes
n = 225
n = 158
n = 99
n = 88
40S
80S
polysome light
polysome heavy
remaining
chaperones
ubiqutitine-mediated proteolysis
proteasome
energy metabolism
proteases
RBPs
E
Pathway enrichment analysis of proteins identified by TS-MS
40S
protein metabolism
nucleic acid metabolism
ribosome biogenesis
RNA-processing
ETC
translation
n = 225
80S
cellular metabolism
RNA-processing
ribosome biogenesis
n = 158
p-value ≤ 0.05
FC ≥ 2
polysome light
cellular metabolism
RNA metabolism
ribosome biogenesis
splicing
translational regulation
gene expression
n = 99
polysome heavy
cellular metabolism
RNA transport
n = 88
F
Distribution of RBPs identified by TS-MS (n=600)
40S 80S polysomes
relative enrichment
G
TS-MS
ARC-MS (all proteins)
1083
488
102
9
12
1
337
centrosomal proteins
TS-MS
ARC-MS (all proteins)
1094
497
103
0
1
0
101
centriolar proteins
TS-MS
ARC-MS (all proteins)
1089
496
103
1
6
0
44
clathrin-coated vesicles

**Fig. S1: ARC-MS faithfully captures translationally active ribosomal complexes**

**(A)** Levels of pluripotency markers and mesoderm markers in hESCs and hESC-derived mesoderm progenitors evaluated by RT-qPCR.

**(B)** Hierarchical clustering of ARC-MS replicates shows a fate specific distinction in proteins recruited on active ribosomes.

**(C)** Heatmap depicting the abundance of 40S and 80S ribosomal proteins in indicated samples.

**(D)** Distribution of proteins detected in indicated ribosomal fractions by TS-MS annotated based of their molecular function depicted as bar graph.

**(E)** Pathway enrichment analysis of proteins from each investigated ribosomal fraction from TS-MS.

**(F)** Heatmap showing the distribution of all detected RBPs in ribosomal fractions detected by TS-MS.

**(G)** ARC-MS allows reliable identification of proteins recruited on active ribosomes. Overlap of proteins that are faithfully identified by ARC-MS, TS-MS to that indicated complexes, predicted as contamination upon sucrose gradient based separation of ribosomal complexes.

Error bars represent ±SEM; p-values calculated using Student’s t-test (n = 3).

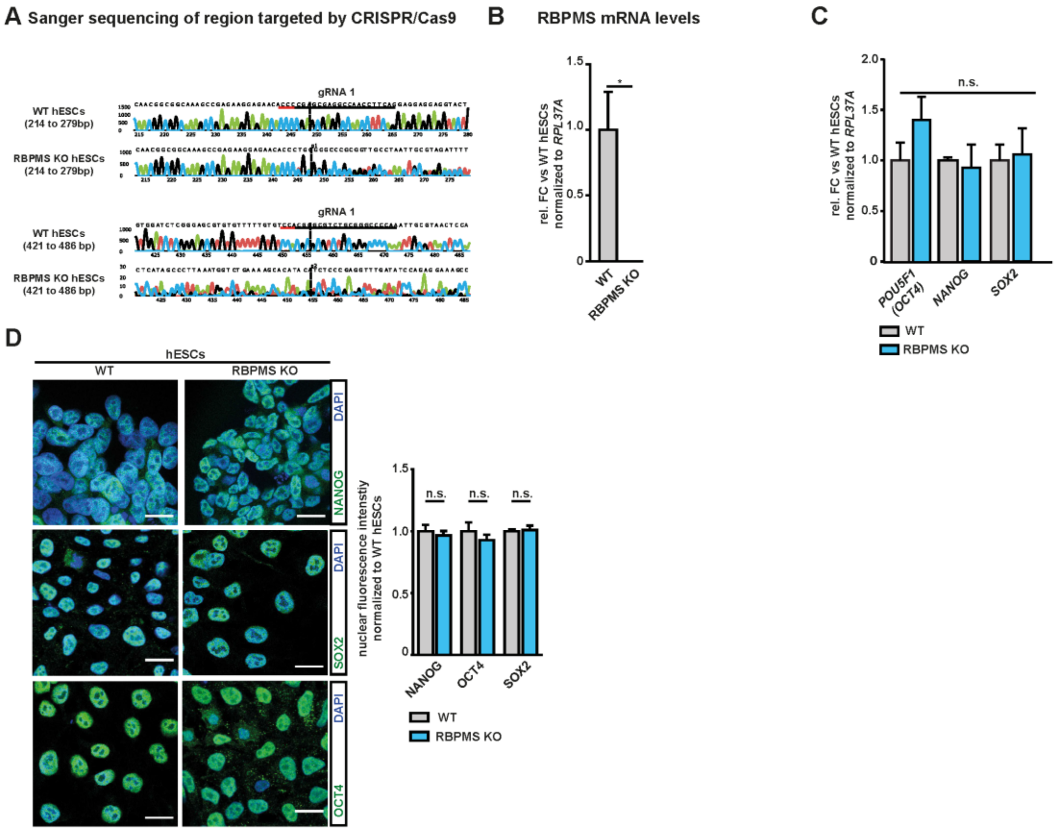
A Sanger sequencing of region targeted by CRISPR/Cas9
gRNA 1
WT hESCs (214 to 279bp)
RBPMS KO hESCs (214 to 279bp)
WT hESCs (421 to 486 bp)
RBPMS KO hESCs (421 to 486 bp)
B RBPMS mRNA levels
rel. FC vs WT hESCs normalized to RPL37A
WT
RBPMS KO
C
n.s.
POU5F1 (OCT4)
NANOG
SOX2
WT
RBPMS KO
D
hESCs
WT
RBPMS KO
DAPI
NANOG
SOX2
OCT4
nuclear fluorescence intenstiy normalized to WT hESCs
n.s.

Figure S2

**Fig. S2: RBPMS loss does not affect the expression of pluripotency factors in hESCs.**

**(A)** Sanger sequencing-based confirmation of genome editing in the generated RBPMS-KO hESCs.

**(B)** Transcript levels of *RBPMS* in RBPMS-KO hESCs compared to WT evaluated by qPCR.

**(C)** Transcript levels of indicated pluripotency factors (*POU5F1* (*OCT4*), *NANOG* and *SOX2*) are not affected by the loss of RBPMS in hESCs, evaluated by RT-qPCR. Bar graphs show relative fold change normalized to *RPL37A*.

**(D)**, RBPMS loss does not affect pluripotency. Representative images of WT and RBPMS-KO hESCs stained for OCT4, SOX2 and NANOG: Bar graph shows normalized expression levels of indicated pluripotency markers.

Error bars represent ±SEM; p-values calculated using Student’s t-test (*≤0.05, **≤0.01, ***≤0.001, ****≤0.0001, n = 3).

**Figure S3**

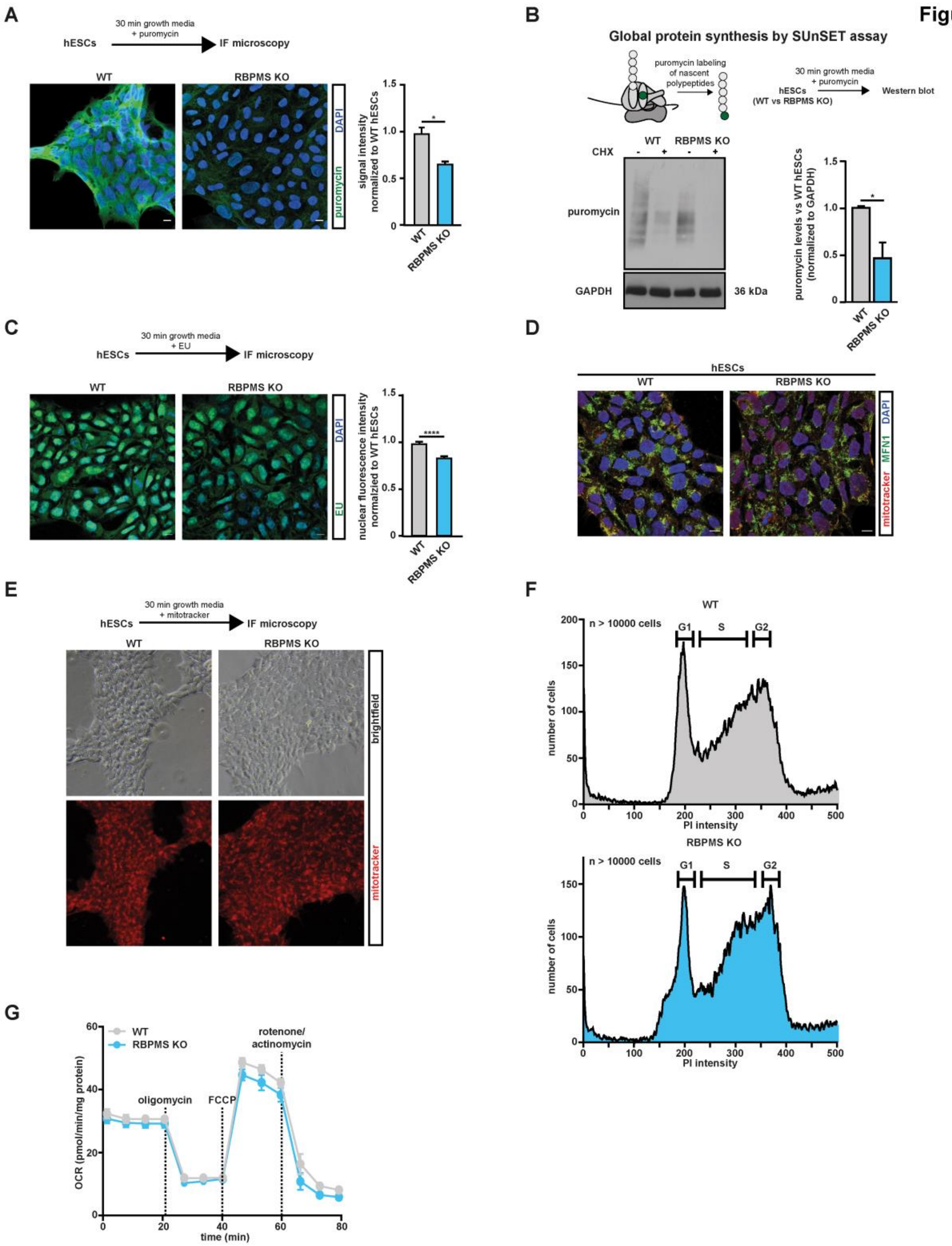
A
30 min growth media
+ puromycin
hESCs
IF microscopy
WT
RBPMS KO
DAPI
puromycin
signal intensity normalized to WT hESCs
B
Global protein synthesis by SUnSET assay
puromycin labeling of nascent polypeptides
30 min growth media
+ puromycin
hESCs
(WT vs RBPMS KO)
Western blot
WT
RBPMS KO
CHX
puromycin
GAPDH
36 kDa
puromycin levels vs WT hESCs (normalized to GAPDH)
C
30 min growth media
+ EU
hESCs
IF microscopy
WT
RBPMS KO
DAPI
EU
nuclear fluorescence intensity normalzied to WT hESCs
D
hESCs
WT
RBPMS KO
DAPI
MFN1
mitotracker
E
30 min growth media
+ mitotracker
hESCs
IF microscopy
WT
RBPMS KO
brightfield
mitotracker
F
WT
n > 10000 cells
G1
S
G2
number of cells
PI intensity
RBPMS KO
n > 10000 cells
G1
S
G2
number of cells
PI intensity
G
WT
RBPMS KO
oligomycin
FCCP
rotenone/
actinomycin
OCR (pmol/min/mg protein)
time (min)

**Fig. S3: RBPMS loss specifically and severely inhibits *de novo* protein synthesis without affecting transcription, mitochondrial morphology, integrity and activity, and cell cycle**

*De novo* protein synthesis evaluated by measuring puromycin in cooperation into nascent polypeptides using anti-puromycin antibody between WT and RBPMS-KO hESCs, evaluated by (**A**) fluorescence microscopy (**B**) by western blotting using anti-puromycin antibody.

(**C**) Representative micrographs depicting the nascent transcription in wild type and RBPMS-KO hESCs, measured by EU (5-Ethynyl Uridine) labeling assay.

Evaluation of **(E)** mitochondrial distribution (mitofusin and mitotracker staining) and **(D)** mitochondrial integrity by live mitotracker intake in RBPMS-KO and WT hESCs.

**(F))** Evaluation of cell cycle upon RBPMS-KO in hESCs**.**

**(G)** Mitochondrial oxygen consumption rates (OCR; pmol $O_2$/min) in WT and RBPMS-KO hESCs. Specific inhibitors used in the analysis are indicated.

Error bars represent ±SEM; p-values calculated using Student's t-test (*≤0.05, **≤0.01, ***≤0.001, ****≤0.0001, n = 3).

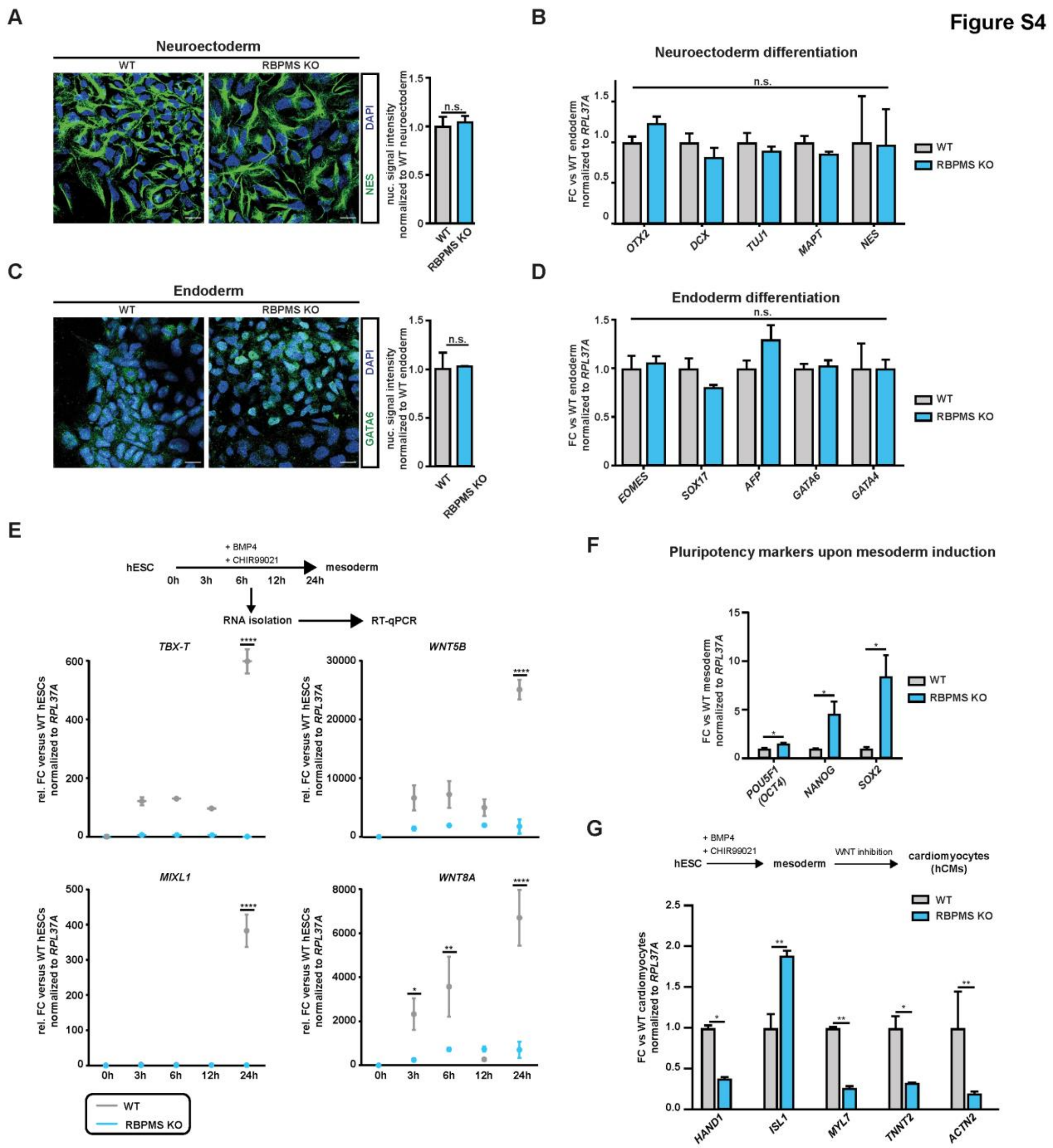
Figure S4
A
Neuroectoderm
WT
RBPMS KO
DAPI
NES
nuc. signal intensity normalized to WT neuroectoderm
n.s.
B
Neuroectoderm differentiation
FC vs WT endoderm normalized to RPL37A
OTX2
DCX
TUJ1
MAPT
NES
C
Endoderm
GATA6
nuc. signal intensity normalized to WT endoderm
D
Endoderm differentiation
EOMES
SOX17
AFP
GATA6
GATA4
E
hESC
+ BMP4
+ CHIR99021
mesoderm
0h 3h 6h 12h 24h
RNA isolation
RT-qPCR
TBX-T
WNT5B
MIXL1
WNT8A
rel. FC versus WT hESCs normalized to RPL37A
F
Pluripotency markers upon mesoderm induction
FC vs WT mesoderm normalized to RPL37A
POU5F1 (OCT4)
NANOG
SOX2
G
WNT inhibition
cardiomyocytes (hCMs)
FC vs WT cardiomyocytes normalized to RPL37A
HAND1
ISL1
MYL7
TNNT2
ACTN2

**Fig. S4: RBPMS loss does not affect the neuroectoderm or endoderm commitment while ablates cardiac mesoderm induction**

Loss of RBPMS, does not affect (**A** & **B)** neuroectoderm and (**C** & **D)** endoderm commitment potential of hESCs, evaluated by immune fluorescence microscopy and qPCR for the indicated markers.

(**E**) Kinetics of induction of mesodermal markers confirms mesoderm commitment defects upon RBPMS loss in hESCs. Plots represent gene expression kinetics of indicated markers normalized to *RPL37A* in RBPMS-KO compared to WT hESCs during mesoderm induction.

**(F)** Loss of RBPMS results in residual expression of indicated pluripotency markers upon mesoderm induction, evaluated by qPCR analysis.

**(G)** Loss of RBPMS impedes terminal differentiation of hESCs to cardiomyocytes indicated by RT-qPCR analysis of cardiomyocyte markers.

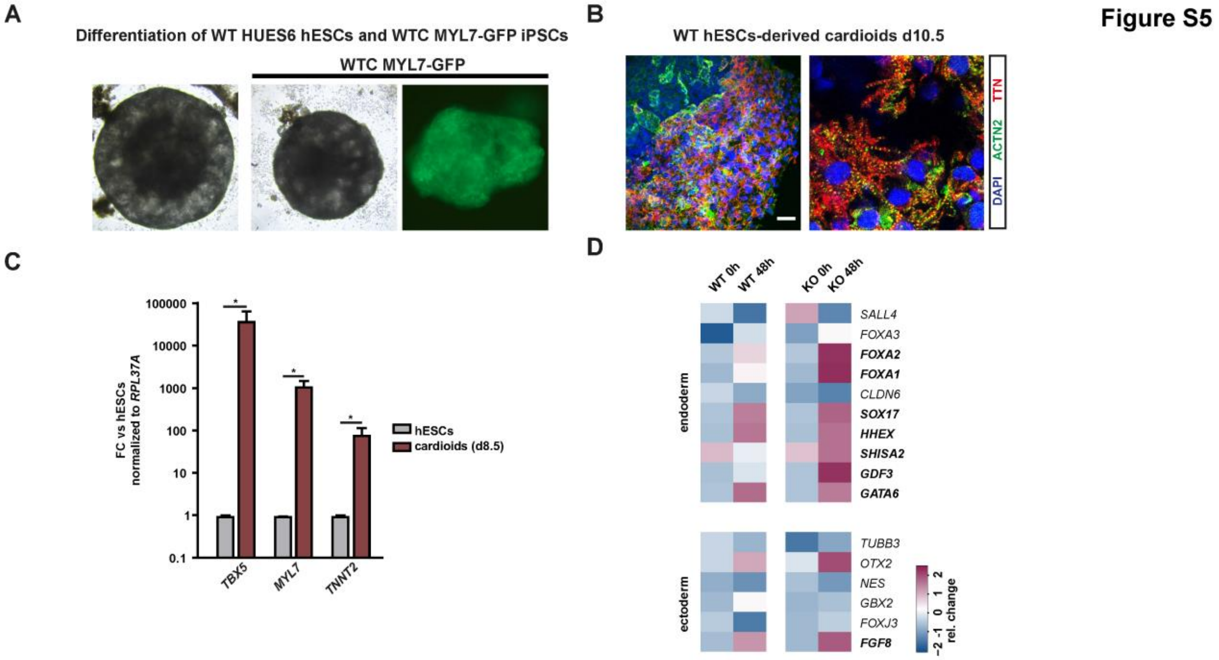

A
Differentiation of WT HUES6 hESCs and WTC MYL7-GFP iPSCs
WTC MYL7-GFP
B
WT hESCs-derived cardioids d10.5
DAPI ACTN2 TTN
Figure S5
C
100000
10000
1000
100
10
1
0.1
FC vs hESCs normalized to RPL37A
hESCs
cardioids (d8.5)
TBX5
MYL7
TNNT2
D
WT 0h
WT 48h
KO 0h
KO 48h
endoderm
SALL4
FOXA3
FOXA2
FOXA1
CLDN6
SOX17
HHEX
SHISA2
GDF3
GATA6
ectoderm
TUBB3
OTX2
NES
GBX2
FOXJ3
FGF8
rel. change
−2 -1 0 1 2

**Fig. S5: Cardioids faithfully recapitulate human cardiac commitment**

**(A)** The cardioid induction protocol yields cardiac organoids with typical morphology and expression of cardiomyocytes markers, confirmed using hiPSC-based cardiac reporter line, *MYL7-GFP*.

**(B)** Immunostainings for cardiac-specific ACTN2 and TTN for d10.5 cardioids derived from HUES6 hESCs (WT hESCs).

**(C)** RT-qPCR showing the expression of indicated cardiomyocyte marker genes in WT hESC-derived cardiods.

**(D)** Heatmaps showing expression of endoderm and ectoderm markers after mesoderm induction in RBPMS-KO hESCs w.r.t WT hESCs.

### Figure S6

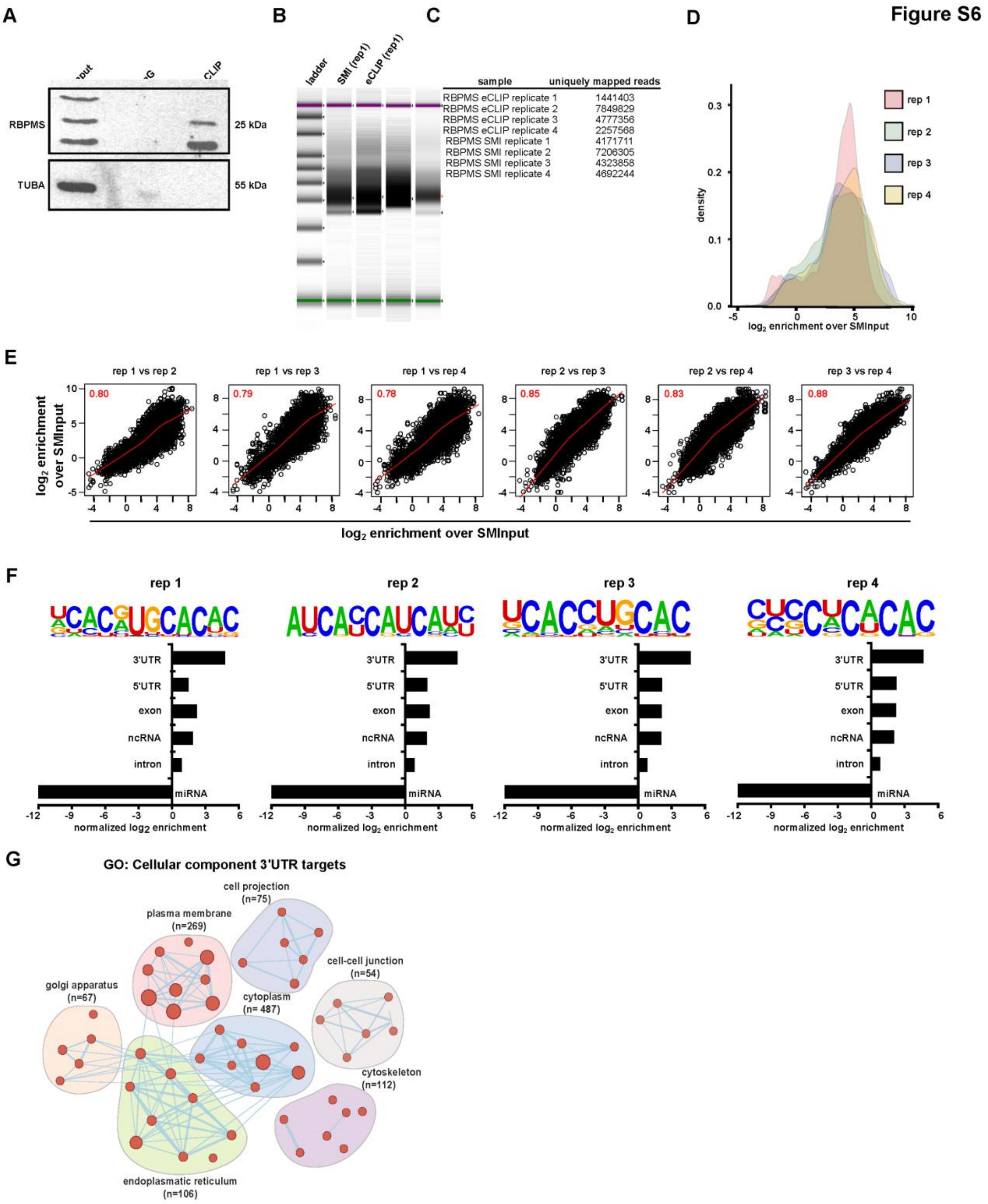

A
B
C
D
RBPMS
TUBA
25 kDa
55 kDa
CLIP
ladder
SMI (rep1)
eCLIP (rep1)
sample
uniquely mapped reads
RBPMS eCLIP replicate 1 1441403
RBPMS eCLIP replicate 2 7849829
RBPMS eCLIP replicate 3 4777356
RBPMS eCLIP replicate 4 2257568
RBPMS SMI replicate 1 4171711
RBPMS SMI replicate 2 7206305
RBPMS SMI replicate 3 4323858
RBPMS SMI replicate 4 4692244
rep 1
rep 2
rep 3
rep 4
density
log2 enrichment over SMInput
E
rep 1 vs rep 2
rep 1 vs rep 3
rep 1 vs rep 4
rep 2 vs rep 3
rep 2 vs rep 4
rep 3 vs rep 4
0.80
0.79
0.78
0.85
0.83
0.88
log2 enrichment over SMInput
F
3'UTR
5'UTR
exon
ncRNA
intron
miRNA
normalized log2 enrichment
G
GO: Cellular component 3'UTR targets
cell projection (n=75)
plasma membrane (n=269)
cell-cell junction (n=54)
golgi apparatus (n=67)
cytoplasm (n= 487)
cytoskeleton (n=112)
endoplasmatic reticulum (n=106)

**Fig. S6: eCLIP-seq maps the RBPMS mRNA targets in hESCs.**

**(A)** Representative western blot-based validation of immunoprecipitation performed during eCLIP of RBPMS, TUBA serves as the control.

**(B)** Representative eCLIP libraries, illustrated as tape-station readouts.

**(C)** Uniquely mapped reads for all eCLIP samples.

**(D)** Read density of eCLIP-Seq data indicating fold enrichment over respective SMIinputs.

**(E)** Correlation of enriched eCLIP-peaks over SMInput between each replicate.

**(F)** Top sequence motif significantly bound by RBPMS in the indicated category along with their distribution categorized based their binding on the mRNA coordinates for each replicate.

**(G)** Enriched GO terms (cellular component) found in RBPMS 3'UTR targets.

### Supplementary Figure 7

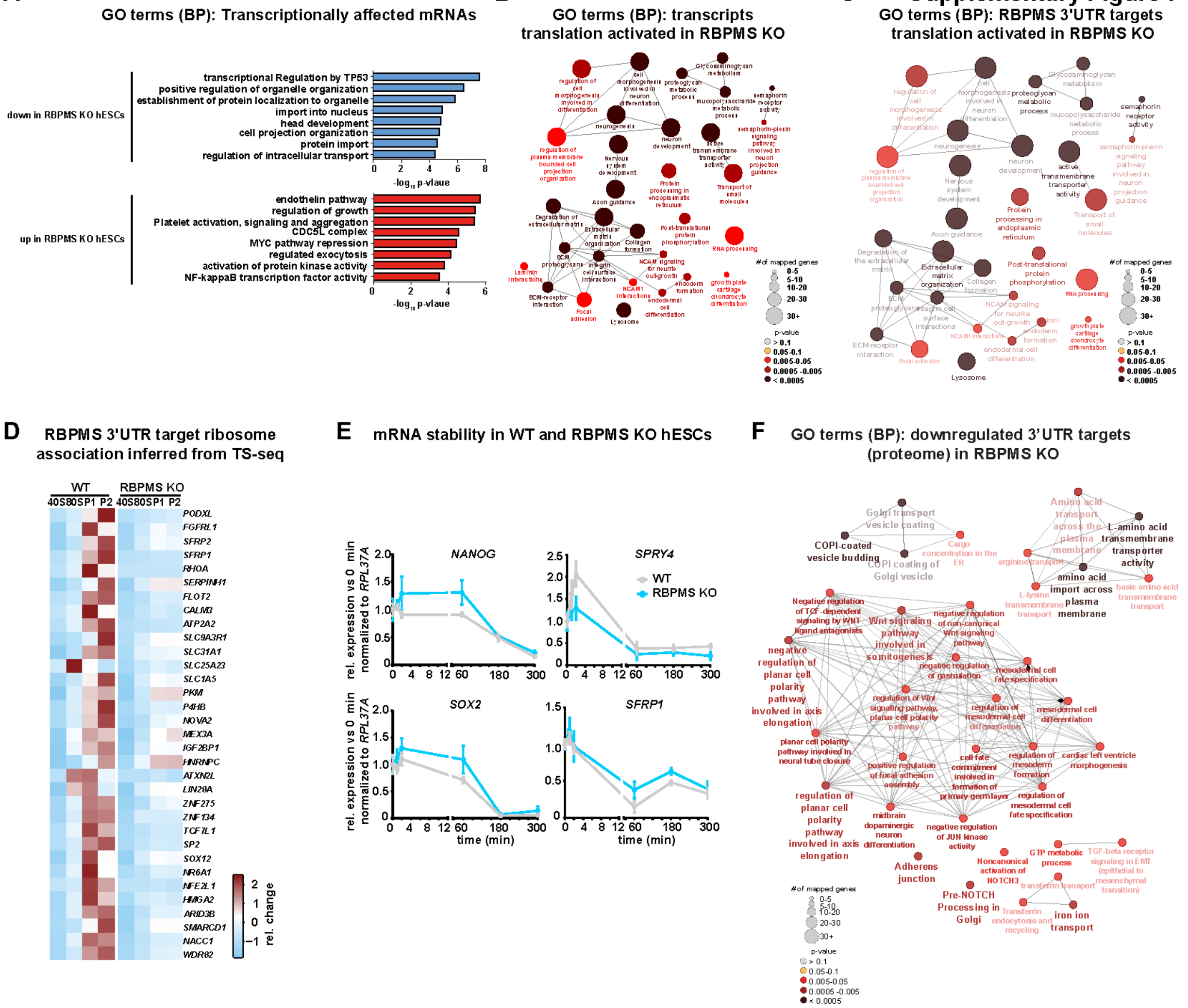

**Fig. S7: RBPMS loss leads to depletion of proteins encoded by its target mRNAs without affecting mRNA abundance**

**(A)** Functional annotation of genes affected only at the level of transcription (changing only at mRNA level), upon loss of RBPMS, does not suffice for the molecular and developmental defects due to the absence of RBPMS in hESCs.

**(B)** Gene Ontology-based functional enrichment analysis for translationally activated transcripts in RBPMS-KO hESCs (significance level indicated by color code).

**(C)** Gene Ontology-based functional enrichment analysis for 3'UTR targets translationally activated in RBPMS-KO hESCs (significance level indicated by color code).

**(D)** Heatmap of ribosome occupancy of selected RBPMS 3'UTR targets in WT hESCs and RBPMS-KO hESCs.

**(E)** Loss of RBPMS does not change the stability of indicated mRNAs, including RBPMS 3'UTR targets, revealed by time-course experiment upon Actinomycin B treatment.

**(F)** Gene ontology analysis of 3'UTR target proteins depleted in RBPMS-KO hESCs w.r.t WT hESCs (significance levels indicated in the legend depicted at the right).

Error bars represent ±SEM; p-values calculated using Student's t-test (n=3).

**Figure S8**

**A**

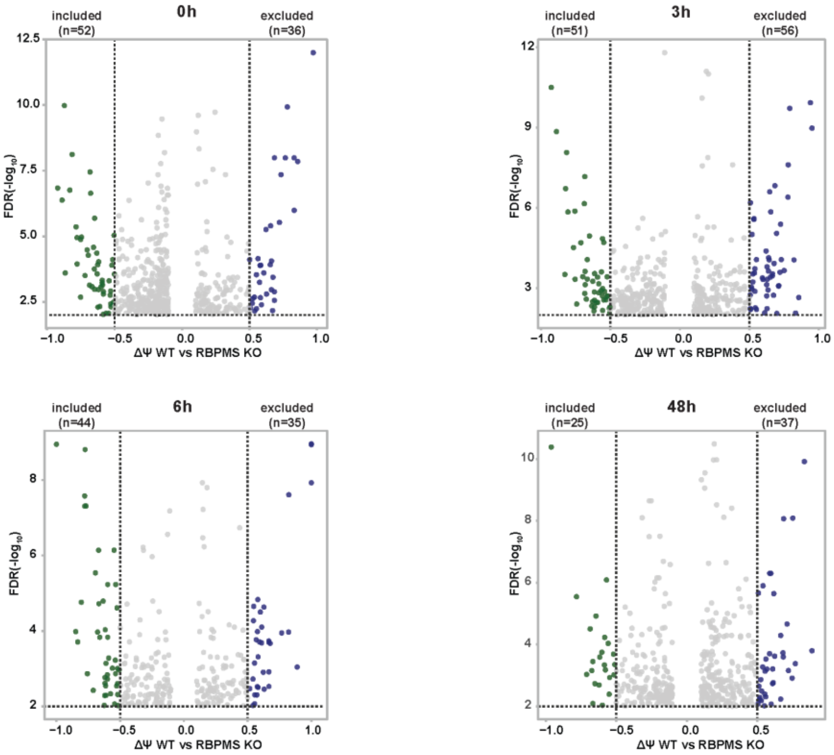


**B**

RBPMS eCLIP targets

| | 0h | | | | | 3h | | | |
|---|---|---|---|---|---|---|---|---|---|
| | 3'UTR | 5'UTR | intron | exon | | 3'UTR | 5'UTR | intron | exon |
| included | 0 | 0 | 2 | 1 | included | 3 | 0 | 1 | 0 |
| excluded | 0 | 0 | 0 | 0 | excluded | 0 | 0 | 3 | 0 |

| | 6h | | | | | 48h | | | |
|---|---|---|---|---|---|---|---|---|---|
| | 3'UTR | 5'UTR | intron | exon | | 3'UTR | 5'UTR | intron | exon |
| included | 1 | 0 | 2 | 0 | included | 0 | 0 | 2 | 0 |
| excluded | 0 | 0 | 1 | 0 | excluded | 3 | 0 | 3 | 1 |

**Fig. S8: RBPMS loss does not affect pre-mRNA splicing in pluripotency and during cardiac mesoderm commitment**

**(A)** Effect of the loss of RBPMS on global changes on pre-mRNA splicing in pluripotency and upon mesoderm commitment (0h (hESCs), 3h, 6h and 48h (mesoderm) into mesoderm induction) depicted as volcano plot of $\Delta\psi$ and FDR (dotted lines indicate significance thresholds FDR $\leq$ 0.01 and $\Delta\psi \leq -0.5$ or $\geq 0.5$).

**(B)** Table summarizing splicing changes for RBPMS eCLIP-targets (3'UTR, 5'UTR, intron and exon) at indicated timepoints during mesoderm induction.

Error bars represent ±SEM; p-values calculated using Student's t-test (n.s.>0.5, *≤0.05, **≤0.01, ***≤0.001, ****≤0.0001, n = 3).

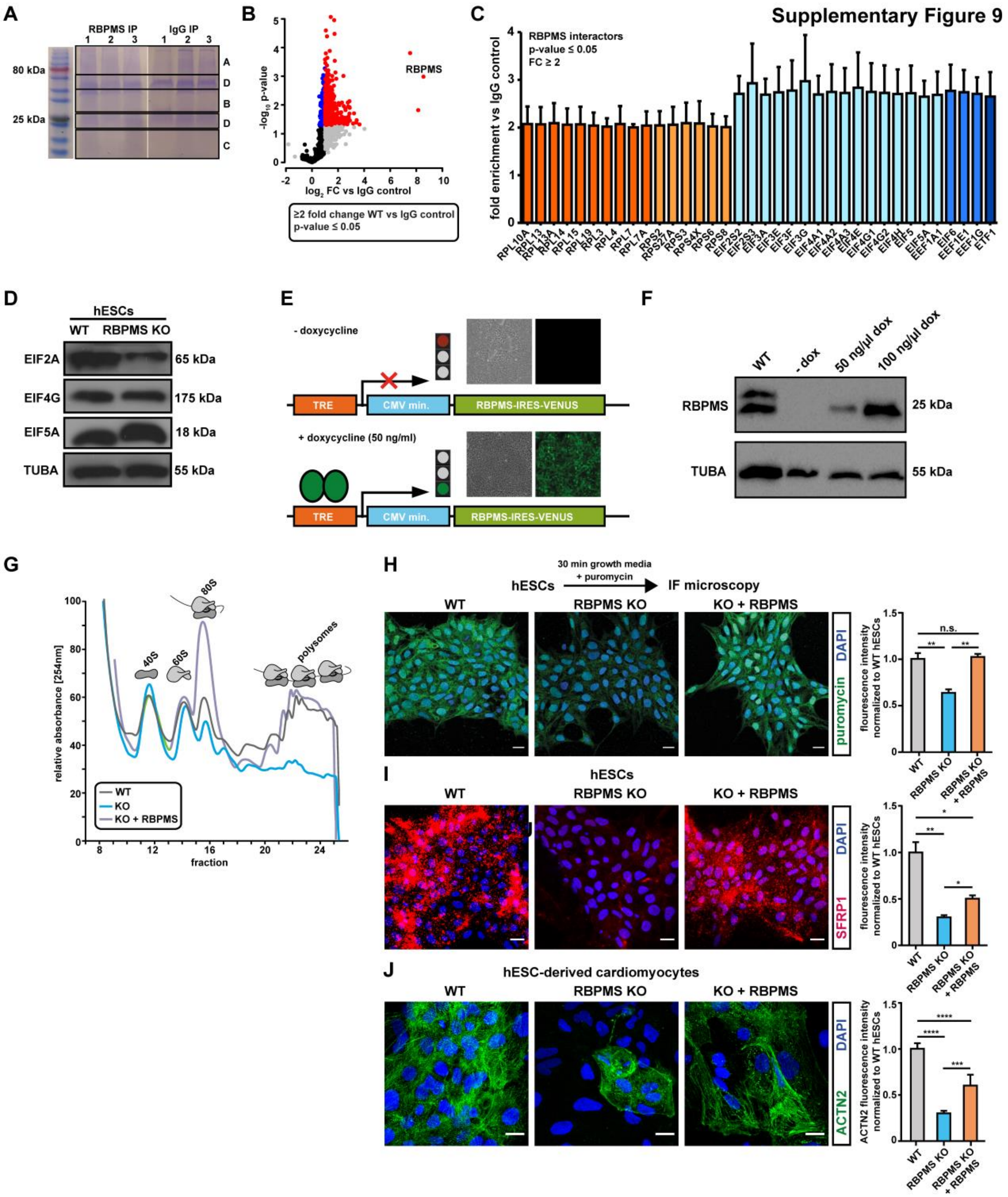
Supplementary Figure 9
A
RBPMS IP
IgG IP
80 kDa
25 kDa
B
RBPMS
-log10 p-value
log2 FC vs IgG control
≥2 fold change WT vs IgG control
p-value ≤ 0.05
C
RBPMS interactors
p-value ≤ 0.05
FC ≥ 2
fold enrichment vs IgG control
D
hESCs
WT
RBPMS KO
EIF2A
EIF4G
EIF5A
TUBA
65 kDa
175 kDa
18 kDa
55 kDa
E
- doxycycline
+ doxycycline (50 ng/ml)
TRE
CMV min.
RBPMS-IRES-VENUS
F
WT
- dox
50 ng/µl dox
100 ng/µl dox
RBPMS
TUBA
25 kDa
55 kDa
G
relative absorbance [254nm]
fraction
40S
60S
80S
polysomes
WT
KO
KO + RBPMS
H
30 min growth media + puromycin
hESCs
IF microscopy
WT
RBPMS KO
KO + RBPMS
DAPI
puromycin
flourescence intensity normalized to WT hESCs
I
hESCs
SFRP1
J
hESC-derived cardiomyocytes
ACTN2
ACTN2 fluorescence intensity normalized to WT hESCs

**Fig. S9: Controlled, timely re-expression of RBPMS rescues the molecular, cellular and development defects caused by RBPMS deficiency in hESCs**

**(A)** Coomassie blue-stained gels depicting the indicated eluates from RBPMS pull down. Size separated proteins were pooled independently, isotope-labeled, and quantified.

**(B)** Volcano plot showing specific enrichment of proteins upon RBPMS pull-down w.r.t IgG control, in triplicates.

**(C)** Specific enrichment of ribosomal proteins and translation factors upon RBPMS pull down.

**(D)** Total levels of the indicated proteins in RBPMS KO hESCs compared to WT.

**(E)**, **(F)** Timely reconstitution of RBPMS in RBPMS-KO hESCs rescues (G)**,** ribosome occupancy defects, (**H)** general protein synthesis defects, (**I**) translation defect of representative 3'UTR target of RBPMS as well as (**J)** lineage commitment defects.

Error bars represent ±SEM; p-values calculated using Student's t-test (n = 3).
